# Phylogenetics, Trait Covariance Analysis, and the Evolution of Fin and Body Shape in the Surgeonfishes

**DOI:** 10.1101/2025.10.09.680739

**Authors:** Linnea L. Lungstrom, Mireille Farjo, Ryan Isdonas, Andrew B. George, Mark W. Westneat

## Abstract

Insights into ecomorphology are enriched when investigating evolutionary correlations between ecologically and functionally important traits. However, tools currently available to model patterns of shape covariation limit researchers to either assume that trait covariances between taxa are independent of, or completely described by, their shared evolutionary history. Using a novel Pagel’s λ adjustment to incorporate phylogenetic signal in trait covariance analysis, we aim to solve this long-standing problem in phylogenetic comparative methods and investigate ecological associations and evolutionary shape covariation in the ecologically and morphologically diverse case study of surgeonfishes (Acanthuridae). By revising acanthurid phylogenetic relationships and analyzing the geometric morphometrics of their body, head, and fins, we found that head and body shape were significantly associated with dietary ecotype. Surgeonfishes showed a significant negative correlation between caudal fin and pectoral fin shape; high/low aspect ratio (AR) tails are associated with low/high AR pectoral fins, respectively, suggesting locomotor tradeoffs. With our new approach to estimate the influence of phylogeny on trait covariances between taxa, we found the caudal fin covaried with both the body and pectoral fin due to dietary and locomotor demands, respectively and exhibited the highest evolutionary variance along the primary axis of integration in all trait covariance comparisons.

## Introduction

Ecomorphological studies provide critical insight into how the interaction between morphology and ecology relates to the adaptive diversification and survival of species. While many ecomorphological studies focus on how individual morphologies relate to ecology (Friedman et al., 2016, 2019; Tavera et al., 2018), functional performance (Chang et al., 2012; Collar et al., 2009; Friedman et al., 2021; Krishnadas et al., 2018; Langerhans, 2008; Sambilay, 1990; Walker et al., 2013; Walker & Westneat, 2002; Webb, 1982, 1984; Weihs, 1973; Westneat, 1996), and evolutionary rates (Du et al., 2019; Price et al., 2019), work has also examined how shape variation in key traits may covary with one another in the context of morphological, ecological, or functional diversification (Aguilar-Medrano et al., 2016; Feilich, 2016; Goswami & Polly, 2010; Larouche et al., 2015, 2018). While shape covariation has been shown to both constrain and facilitate adaptive diversification (Du et al., 2019; Evans et al., 2017; Goswami et al., 2014; Goswami & Polly, 2010; Klingenberg, 2013; Wagner & Altenberg, 1996), questions still remain. For example, are certain forms more correlated than others across an organism? Does the degree of correlation between shapes relate to an organism’s ecology or biomechanics? To what extent do the shared evolutionary histories of species explain patterns of covariation between shapes, and are the evolutionary variances along the primary axes of integration different or similar depending on the strength of association between shapes?

Ecomorphology of marine fishes is an ideal system in which to ask these questions. Head shape and body elongation are functionally important traits for fishes as they are associated with habitat (especially along the benthic-pelagic axis), trophic niches, and feeding modes (Bejarano et al., 2017; Claverie & Wainwright, 2014; Cooper & Westneat, 2009; Floeter et al., 2018; Friedman et al., 2016, 2019; López-Fernández et al., 2013; Mahe et al., 2014; Robertson et al., 1979; Siqueira et al., 2020; Tavera et al., 2018). For example, smaller mouths and deeper bodies have been attributed to benthic-associated fishes (Claverie & Wainwright, 2014). Habitat use is also linked to functional swimming modes (Fulton et al., 2001) that are facilitated by specific body forms (Collar et al., 2009; Friedman et al., 2021; Walker et al., 2013) and fin shapes (Walker & Westneat, 2002; Webb, 1984; Westneat, 1996). For example, the aspect ratio (AR, a measure of paddle vs wing-like shape) of pectoral and caudal fins in fishes have been linked to locomotor efficiency, thrust, speed, endurance, and maneuverability trade-offs (Chang et al., 2012; Gerstner, 1999; Krishnadas et al., 2018; Sambilay, 1990; Walker & Westneat, 2000, 2002; Webb, 1982; Weihs, 1973; Westneat et al., 2017). Body (Price et al., 2019) and fin (Du et al., 2019) shapes have been shown to evolve at different rates across fishes, with locomotor mode and dietary ecotype often significantly associated with selective evolutionary models (Friedman et al., 2016; McCord et al., 2021; Ribeiro et al., 2018, Santaquiteria et al., 2026). Morphological integration is also related to rapid diversification in fishes (Burns et al., 2023; Evans et al., 2021; Larouche et al., 2018), highlighting the importance of evolutionary covariation among traits in fishes.

A long-standing problem in phylogenetic comparative methods limits the ability to answer questions about evolutionary shape covariation. Currently, there are two ways researchers can quantify morphological covariation between shapes using two-block partial least squares (2B-PLS) models (Adams et al., 2025; Baken et al., 2021). First, these models can assume the evolution of the multivariate trait data is not influenced by taxa’s shared evolutionary history (functions “two.b.pls” and “integration.test” in R package “geomorph”, (Adams et al., 2025; Baken et al., 2021). Alternatively, these models can assume the evolution of the multivariate trait data is entirely consistent with the phylogeny, under a Brownian motion (BM) model of evolution (“phylo.integration” function in “geomorph”, Adams et al., 2025; Adams & Felice, 2014; Baken et al., 2021). This approach assumes that trait covariances between taxa are completely described by their shared evolutionary history. How do researchers deal with systems in which the evolution of the multivariate trait data is neither independent of, nor completely explained by, phylogenetic relationships? Although we can estimate the phylogenetic influence in traits themselves, new tools are needed to model the degree to which shared phylogenetic history influences evolutionary covariances between shapes. Few biological systems can be thought in terms of an “all or nothing” influence of phylogeny on trait covariances, especially those in which selection is acting on shape (Monteiroa, 2013; Revell, 2010). If the true evolutionary process (selection, constraint, time varying rates, etc.) results in a deviation from this “all or nothing” expectation, then the current tools may result in biased estimates of shape correlation. Here, we propose a solution to this long-standing issue in phylogenetic comparative methods by making use of Pagel’s λ, a scalar of the phylogenetic covariances that is evolutionary mode-agnostic, to estimate the influence of shared evolutionary history on shape covariation within the surgeonfishes (Acanthuridae).

Surgeonfishes are an ideal family of fishes in which to investigate ecomorphological associations and implement our new approach to shape covariation analyses. Variation in their head, body, and fin shape has been linked to specific dietary ecologies (Claverie & Wainwright, 2014; Friedman et al., 2016). Surgeonfishes also swim in diverse ways, using primarily either their caudal fin or their pectoral fins to swim (Fulton, 2007; Fulton et al., 2005), potentially leading to shape covariations within the family. Dietary ecotypes differ in both diet and microhabitat, ranging from benthic herbivores grazing on algae, to pelagic planktivores, with some species exhibiting omnivory (Clements et al., 2003; Winterbottom, 1971). Thus, maintaining these different dietary ecotypes may involve specific, and integration of, head, body, and fin morphologies. Additionally, ecomorphological evolution is consistent with adaptive dynamics (Friedman et al., 2016), not BM, in surgeonfishes, suggesting they may deviate from the “all or nothing” assumption of phylogenetic influence on multivariate trait data. Thus, the central aims of this study are to 1) provide a new well-resolved phylogeny for the surgeonfishes as a framework for ecomorphological and evolutionary covariance analyses, 2) determine if head, body, or fin morphology are related to dietary ecotype, 3) investigate the degree of morphological covariation among these subsets by presenting a Pagel’s λ adjustment as a solution to a long-standing problem in phylogenetic comparative methods, and 4) examine how fishes’ ecology or shared evolutionary history may relate to the strength of shape correlation between functionally important traits within the surgeonfishes.

## Materials and Methods

### Time-calibrated phylogenetic tree

To develop a new, well-resolved surgeonfish phylogeny, DNA sequences for 125 fish species were analyzed, including 80 species of surgeonfishes and 45 outgroup taxa from several closely related fish families (Supplementary Appendix A). Sequences for each gene were aligned in Mesquite (Maddison & Maddison, 2025) and imported into IQ-TREE 2 (Kalyaanamoorthy et al., 2017; Minh et al., 2020) to construct the best fit partitioning scheme (Table S1) and a maximum likelihood tree for Bayesian analyses (details in Supplementary Methods). To obtain a time-calibrated phylogenetic hypothesis of Acanthuridae, we estimated divergence times using an optimised relaxed clock (ORC) model with two fossil calibrations and used a standard birth death rate model as a tree prior (details in the Supplementary Methods). We ran two independent analyses for 200 million generations, sampling every 5000 generations in BEAST 2.7.7 (Bouckaert et al., 2019; Drummond et al., 2002, 2006). Convergence between the two runs and effective sample sizes (ESSs) of all model parameters were used to ensure sufficient independent samples using Tracer v.1.7.2 (Rambaut et al., 2018). A single time calibrated tree was generated from the posterior distribution of trees (first 50% removed as burn-in) using LogCombiner 2.6.3 to combine the trees. Subsequently, TreeAnnotator 2.6.3 (Drummond & Rambaut, 2007) was used to produce the maximum clade credibility tree annotated with mean/median branch lengths and to calculate posterior probabilities for node support.

### Geometric morphometrics

To describe the variation in shape across 80 acanthurid species, we quantified head, body, and fin morphologies across 230 2D lateral specimen photographs using geometric morphometrics (details in Supplementary Methods, Supplementary Appendix B). Using the R (version 4.4.2, R Core Team, 2024) package “StereoMorph” (Olsen & Westneat, 2015), we placed a total of 183 landmarks (34 fixed landmarks and 149 sliding semilandmarks along 11 curves) on each specimen image (Figure 1, Tables S2 & S3), capturing the variation in body, head, and fin shapes. Orientation of soft, movable anatomical elements such as fins, and variable features such as jaw angle, require standardization of position across images to avoid including non-biologically relevant axes of variation into the datasets. We developed a Mac app called SurgeonShape (Supplementary Methods; Figure S1; https://github.com/mwestneat/SurgeonShape) to spread and/or rotate fins and jaws to standard positions as well as to calculate comparative and functional traits of the surgeonfish body and fins, including pectoral and caudal fin ARs and body elongation ratio (Figure 1, details in the Supplementary Methods).

**Figure 1.**
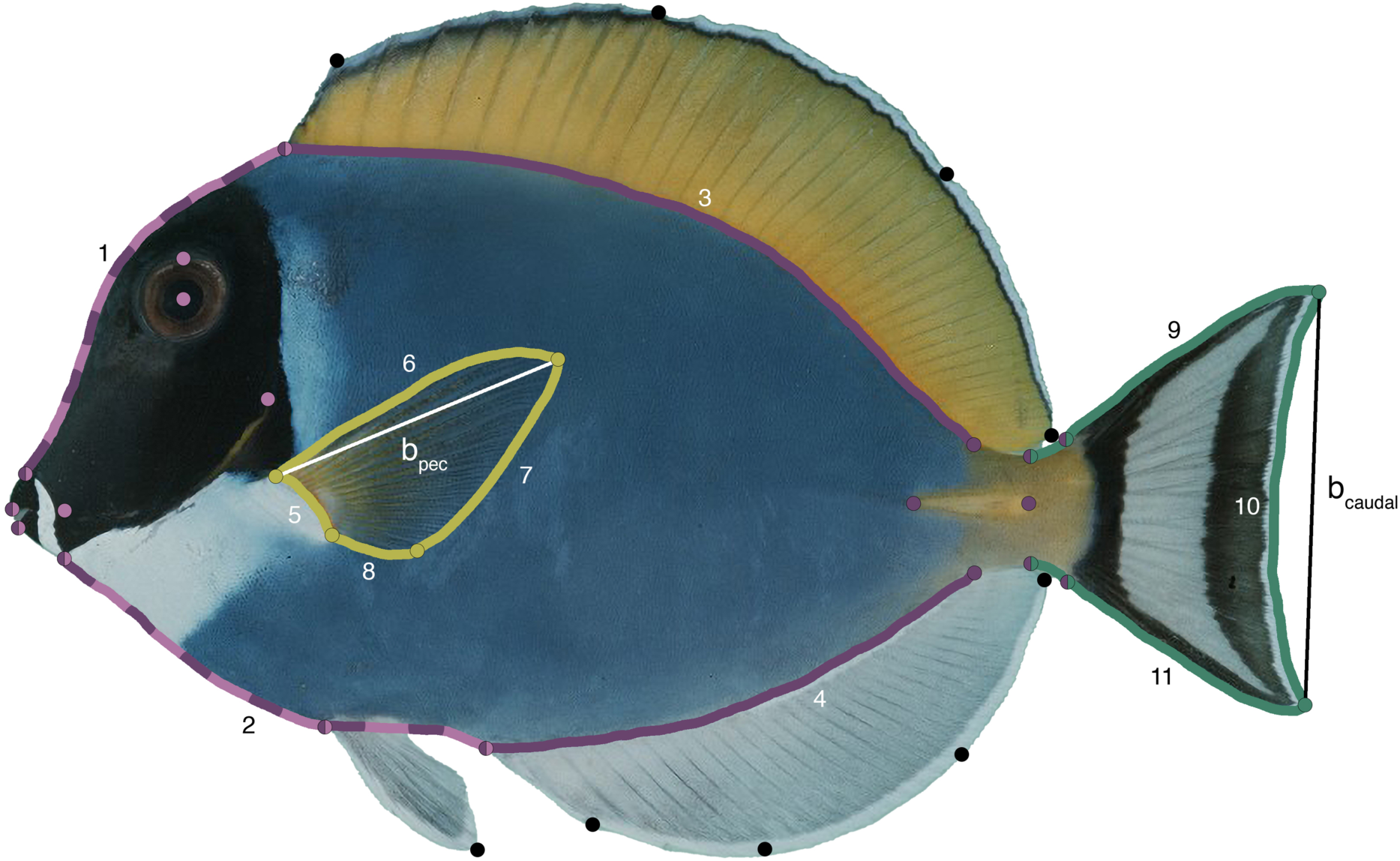
Geometric Morphometric Landmark Scheme. Circles represent fixed landmarks, while numbered curves represent sliding semilandmarks. Circles and curves are color-coded based on which landmark subset they belong to: pink denotes head, purple denotes body, yellow denotes pectoral fin, and green denotes caudal fin. Black circles denote landmarks that are not found in any subset but are included in the overall shape of the fish. Lines b_pec_ and b_caudal_ represent the span of the pectoral and caudal fin used in the respective fin AR calculations.

The visually leveled, rotated, and standardized landmarks were then imported into R and divided into four subsets: caudal fin, pectoral fin, head, and body (Figure 1, Supplementary Methods). For each landmark subset, we grouped digitized images by species, and then projected their landmarks into linear tangent space by performing a generalized Procrustes analysis (GPA) using the “gpagen” function in “geomorph” (Adams et al., 2025; Baken et al., 2021) to remove any non-shape differences in landmark position attributed to differences in rotation, translation, and scaling. Semilandmarks were slid to minimize bending energy. We calculated species mean shapes for each species-group using the “mshape” function in “geomorph” (Adams et al., 2025; Baken et al., 2021) and then used the “findMeanSpec” function in “geomorph” (Adams et al., 2025; Baken et al., 2021) to determine which image was closest to the mean shape for each species. This image was chosen as the representative of that species. When there were only two images of a particular species (which results in a “tie” with similarity to the mean shape), we chose the image that had the least distortion. With a single representative image for each species, we performed a GPA on the raw coordinates that optimized semilandmark positions by reducing bending energy and projected Procrustes-aligned shape data in a linear tangent space using the “gpagen” function in “geomorph” (Adams et al., 2025; Baken et al., 2021) for each subset. We down-sampled the semilandmarks from each landmark subset (Supplementary Methods) and then performed a principal component analysis (PCA) using the “gm.prcomp” function in “geomorph” (Adams et al., 2025; Baken et al., 2021) to visualize the distribution of taxa in morphospace. For each subset, we determined outliers using the “plotOutliers” function in “geomorph” (Adams et al., 2025; Baken et al., 2021), removed these taxa from the dataset, and pruned the tree using the function “drop.tip” in “ape” (Paradis & Schliep, 2019) to exclude outliers and outgroups. Phylomorphospaces were then generated by projecting the pruned time-calibrated phylogeny onto the morphospace to assess evolutionary directionality of shape changes across Acanthuridae.

### Dietary ecotype categorization

Through an extensive literature review (Supplementary Appendix C), we categorized surgeonfishes’ dietary ecotype to investigate our ecomorphological aims. While previous studies (see Supplementary Appendix C) use the planktivore/herbivore dichotomy to bin species, we categorized surgeonfishes into three ecotypes: benthic herbivores/detritivores, pelagic planktivores, and omnivores. Species were considered omnivorous if explicitly stated in the literature as being omnivorous, or if their primary diet was accompanied by multiple accounts of feeding on secondary prey types.

### Statistical analyses

We assessed the relationships between surgeonfish morphology and dietary ecotype by performing a permutations-based phylogenetic Procrustes ANOVA on the Procrustes shape variables using the “procD.pgls” function in “geomorph” (Adams et al., 2025; Baken et al., 2021; Collyer & Adams, 2018, 2024) with model structure of coords ∼ diet. We used residual randomization in a permutation procedure (RRPP) and performed 1000 iterations for significance testing. This analysis assumes that phylogenetic covariances among species perfectly explain residual error in the model (Collyer & Adams, 2018, 2024). If invalid, this assumption has been shown to lead to increased variance in parameter estimates, reduced power, and, in some cases, elevated type I error rates in phylogenetic regressions (Monteiroa, 2013; Revell, 2010). Thus, estimating the amount of phylogenetic signal in the error of our model is critical, especially considering surgeonfish linear morphological traits likely evolved under a model with selection towards an optimum, not BM (Friedman et al., 2016). We estimated the amount of phylogenetic signal (Pagel’s λ) in the residual error of our Procrustes ANOVA models using methods outlined in previous studies (Natale & Slater, 2022; Navalón et al., 2019). In brief, we fit a nonphylogenetic Procrustes ANOVA model using the “procD.lm” function in “geomorph” (Adams et al., 2025; Baken et al., 2021; Collyer & Adams, 2018, 2024), extracted the matrix of residuals, and then fit a multivariate Pagel’s λ model to the residuals using the “transformPhylo.ML” function in the R package “motmot” (Puttick et al., 2020). The maximum likelihood estimate of λ derived from this estimation was then used to rescale the phylogeny for our phylogenetic Procrustes ANOVAs using the “rescale” function in the R package “phytools” (Pennell et al., 2014; Revell, 2024). Post hoc tests were conducted using the “pairwise” function in the R package “rrpp” (Collyer & Adams, 2018, 2024) to compare the mean among dietary ecotype groups using a Holm-Bonferroni correction to account for multiple testing.

We also tested whether continuous fin and body metrics were related to each other and dietary ecotype. We determined whether pectoral fin and caudal fin ARs were associated by transforming branch lengths by optimizing λ using maximum likelihood and performing a phylogenetic generalized least-squares (PGLS) analysis using the “pgls” function in “caper” (Orme et al., 2025). Additionally, we used a phylogenetic ANOVA to determine if elongation ratio, pectoral fin AR, and caudal fin AR, were associated with dietary ecotype using the function “phylANOVA” in “phytools” (Pennell et al., 2014; Revell, 2024) with a Holm-Bonferroni correction. Here, we accounted for the phylogenetic signal in our continuous fin and body metrics using the “phylosig” function in “phytools” (Pennell et al., 2014; Revell, 2024) and transformed our phylogeny for analysis using the same function as described in the previous paragraph.

To evaluate the evolutionary covariation between morphological subsets, we performed 2B-PLS tests using the “phylo.integration” function in the R package “geomorph” (Adams et al., 2025; Baken et al., 2021; Collyer & Adams, 2018, 2024) under the current “all or nothing assumptions” as well as our Pagel’s λ adjustment outlined below. We elected not to test associations between head and body subsets, as many landmarks are shared between them. We treated each dataset as a separate block to avoid any covariation due to the variation in their relative size, position, or orientation on the body of the fish.

We estimated the influence of shared evolutionary history on shape covariation using new methodology to appropriately account for phylogenetic signal in a 2B-PLS. Making use of λ, a scalar of the phylogenetic covariances that is evolutionary mode-agnostic, we account for phylogenetic signal both within *and between* blocks simultaneously, instead of assuming that covariation between blocks is entirely explained by the phylogenetic relationship of the taxa. With each 2B-PLS comparison of shapes, we estimated the amount of phylogenetic signal by defining a grid of possible λ values (0 to 1, with increments of 0.01), transforming the phylogeny with each λ value using the “rescale” function in the R package “phytools” (Pennell et al., 2014; Revell, 2024), and then performing 2B-PLS tests with “phylo.integration” in “geomorph” (Adams et al., 2025; Baken et al., 2021) using each transformed phylogeny. For each result, we then computed the multivariate normal likelihood of the paired PLS score vectors, assuming an evolutionary covariance matrix where the diagonal elements represent evolutionary variances of the dominant cross-block latent traits (conditional on the phylogenetic transformation), and the off-diagonals are the evolutionary covariances between the dominant axes for the two blocks (Revell & Harmon, 2008). These off-diagonals can be interpreted as the strength of evolutionary integration. Importantly, r-PLS, the evolutionary correlation between the two blocks, can be obtained directly from this matrix as:

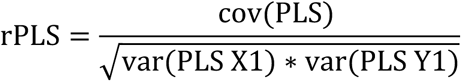

where cov(PLS) is the evolutionary covariation between the two PLS axes and var(PLS X1) and var(PLS Y1) are the evolutionary variances of the first PLS axes for the first and second block respectively.

We chose the λ value that maximized the likelihood of the paired PLS score vectors and used this value to rescale our phylogeny as described above for a final 2B-PLS test. The function “compare.pls” in “geomorph” (Adams et al., 2025; Baken et al., 2021) was used to compare levels of integration between tests. Additionally, for each test, we took the ratio of the evolutionary variances of the PLS1 scores (largest:smallest) to determine how similar blocks were in their evolutionary dispersion along the primary dimension of integration.

## Results

### Acanthurid phylogenetics

Phylogenetic analysis provided a species-rich and well-resolved topology for our comparative analyses. The maximum likelihood tree (Figure S2) and the time-calibrated topology (Figure 2) were nearly identical. Estimated ages and posterior probability support for tree nodes are presented in Figures S3 and S4 respectively. The root node of a monophyletic Acanthuridae was resolved with high support, estimated to be about 54.5 Ma and all but 12 internal nodes within the phylogeny have unambiguous support (Figure 2). Our phylogenetic tree resolves the genus *Naso* as a strongly supported monophyletic group, with its basal divergence at 19.5 Ma, forming the sister group to all other acanthurids. The genus *Prionurus* was also resolved as a monophyletic group with strong support. It was the second group to branch from the backbone topology of the Acanthuridae, approximately 47 Ma, with a crown age of about 16 Ma. The rest of the acanthurids are split into two clades with strong support at 43 Ma, one containing the monotypic genus *Paracanthurus* and genus *Zebrasoma*, and the other containing the genera *Acanthurus* and *Ctenochaetus*. More detailed phylogenetic results can be found in Supplementary Results.

**Figure 2.**
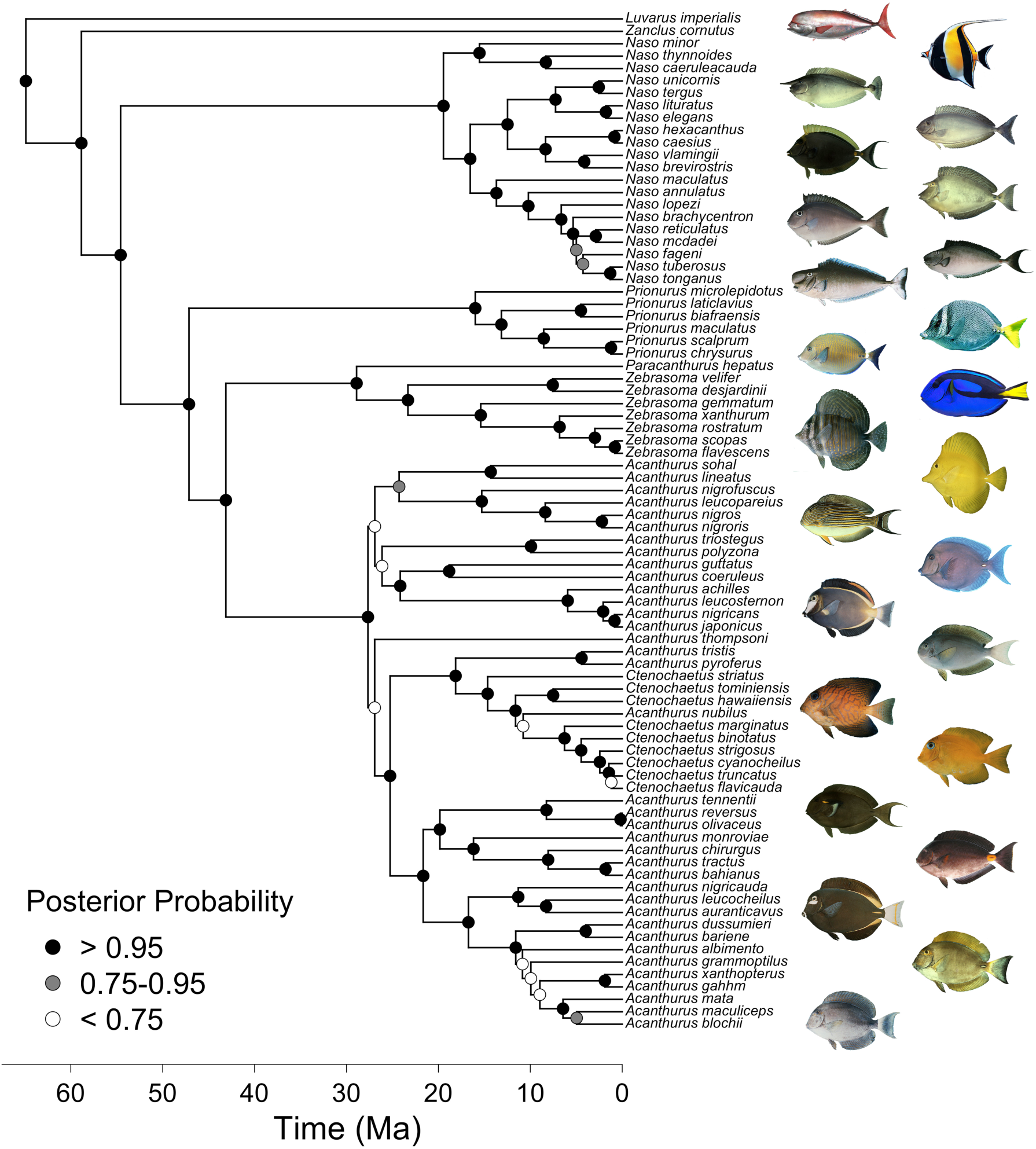
Phylogenetic Tree. Time-calibrated phylogenetic hypothesis for the surgeonfishes, family Acanthuridae, including 80 of 83 surgeonfish species as well as two outgroups (*Zanclus cornutus* and *Luvarus imperialis*) from closely related families. Nodes are color coded by support levels computed as the frequency of occurrence of that node in the post burn-in tree set. Only values above 0.95 represent significant support for that node. Supplementary Materials (Figures S2–S4) provide additional phylogenetic information.

### Elongation and fin aspect ratios

Surgeonfishes exhibit a wide range of elongation ratios and fin ARs. *A. guttatus*, a deeper-bodied species, has the lowest elongation ratio (1.861) while *N. lopezi*, a long-bodied species, has the highest (3.428) (Figure S5, Supplementary Appendix D). When plotted across the phylogeny, we found that members of the genus *Naso* generally exhibit high elongation ratios (apart from *N. literatus*, *N. elegans*, and *N. unicornis*) compared to all other acanthurid species (Figure S5). Pectoral fin ARs range from 1.435 (paddle-like) in *N. maculatus* to 3.102 (wing-like) in *A. sohal* and caudal fin ARs range from 1.251 (paddle-like) in *A. albimento* to 3.878 (wing-like) in *N. thynnoides* (Figure 3, Supplementary Appendix D). Neither caudal nor pectoral fin AR were significantly associated with elongation ratio.

**Figure 3.**
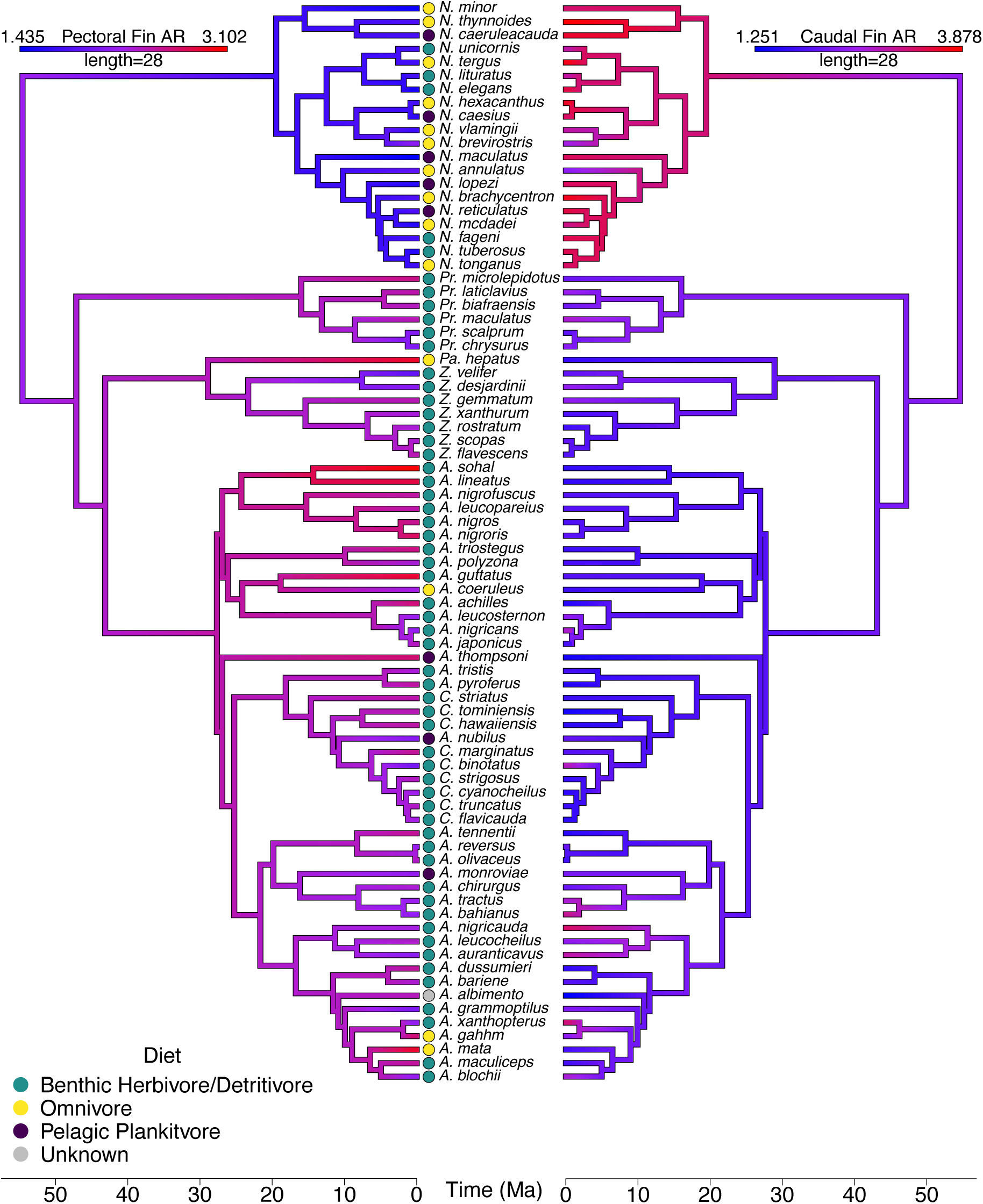
Fin Aspect Ratio Mirror Trees. Caudal and pectoral fin ARs mapped along the Acanthuridae phylogeny by estimating states at internal nodes using maximum likelihood, with caudal fin AR ranging from 1.251 to 3.878 and pectoral fin AR ranging from 1.435 to 3.102. Time is denoted on the bottom scale in Ma. Across the phylogeny, high/low caudal fin AR is generally associated with low/high pectoral fin AR, respectively. Dietary ecotype for each species is represented by colored dots at the tips of trees.

Overall, caudal and pectoral fin AR are significantly negatively correlated even when accounting for shared evolutionary history (λ = 0.458, adjusted R^2^ = 0.284, p < 0.001). Caudal fin AR mapped onto the phylogeny indicates that members of the genus *Naso* generally exhibit high AR caudal fins compared to the rest of the family (Figure 3), although several other *Acanthurus* species demonstrate moderate to high caudal fin ARs. Conversely, an inverse relationship was revealed in pectoral fin AR; *Naso* species exhibited the lowest pectoral fin ARs, with just *Z. velifer* exhibiting a somewhat similar pectoral fin AR. *Ctenochaetus* exhibits an opposite and less extreme relationship to that seen in *Naso*, with many species exhibiting lower caudal fin ARs and higher pectoral fin ARs. This inverse relationship between caudal and pectoral fin AR was not exclusive to *Naso* and *Ctenochaetus*. For example, *P. hepatus*, *A. lineatus*, *A. sohal*, *A. achilles*, *A. guttatus*, *A. thompsoni*, and *A. mata* all exhibit incredibly low caudal fin ARs but some of the highest pectoral fin ARs. Many taxa that exhibit intermediate values of AR in one fin also exhibit intermediate values of AR in the other such as *N. brevirostris*, *Pr. maculatus*, *Pr. chrysurus*, *Z. desjardinii*, *A. leucosternon*, *A. tractus*, *A. nubilus*, *A. monroviae*, *A. leucocheilus*, *A. grammoptilus*, and *A. blochii*. However, some taxa do not follow these trends; *Z. velifer* and *N. annulatus* have low caudal and low pectoral ARs. Although no taxa exhibit both high AR pectoral and caudal fins, *A. nigricauda*, *A. xanthopterus*, and *A. gahhm* have one high AR fin paired with a fin of moderate AR.

### Overall morphological shape patterns

Over 90% of the variation within the complete morphometric dataset is contained in the first nine PCs, with the first two PCs accounting for 70.90% of the total variation in the dataset. PC1 (56.85%) describes a shift from taxa with short, deep bodies and caudal peduncles, flat foreheads with no apex, short caudal fin ends, pointed, wing-like pectoral fins, deep dorsal and anal fins, and small caudal spine areas (negative scores) to taxa with elongated bodies with narrow caudal peduncles, anteriorly peaked, convex foreheads, caudal fins with long trailing edges, round, paddle-like pectoral fins, shallow dorsal and anal fins, and large caudal spine areas (positive scores) (Figure 4). Negative scores on PC2 (14.04%) correspond to taxa with convex foreheads, concave caudal fins with long training edges, and pointed pectoral fins with positive scores corresponding to taxa with flat foreheads, convex caudal fins, and rounder pectoral fins. These differences described by the first two PC axes nearly separate genera into distinct regions of morphospace.

**Figure 4.**
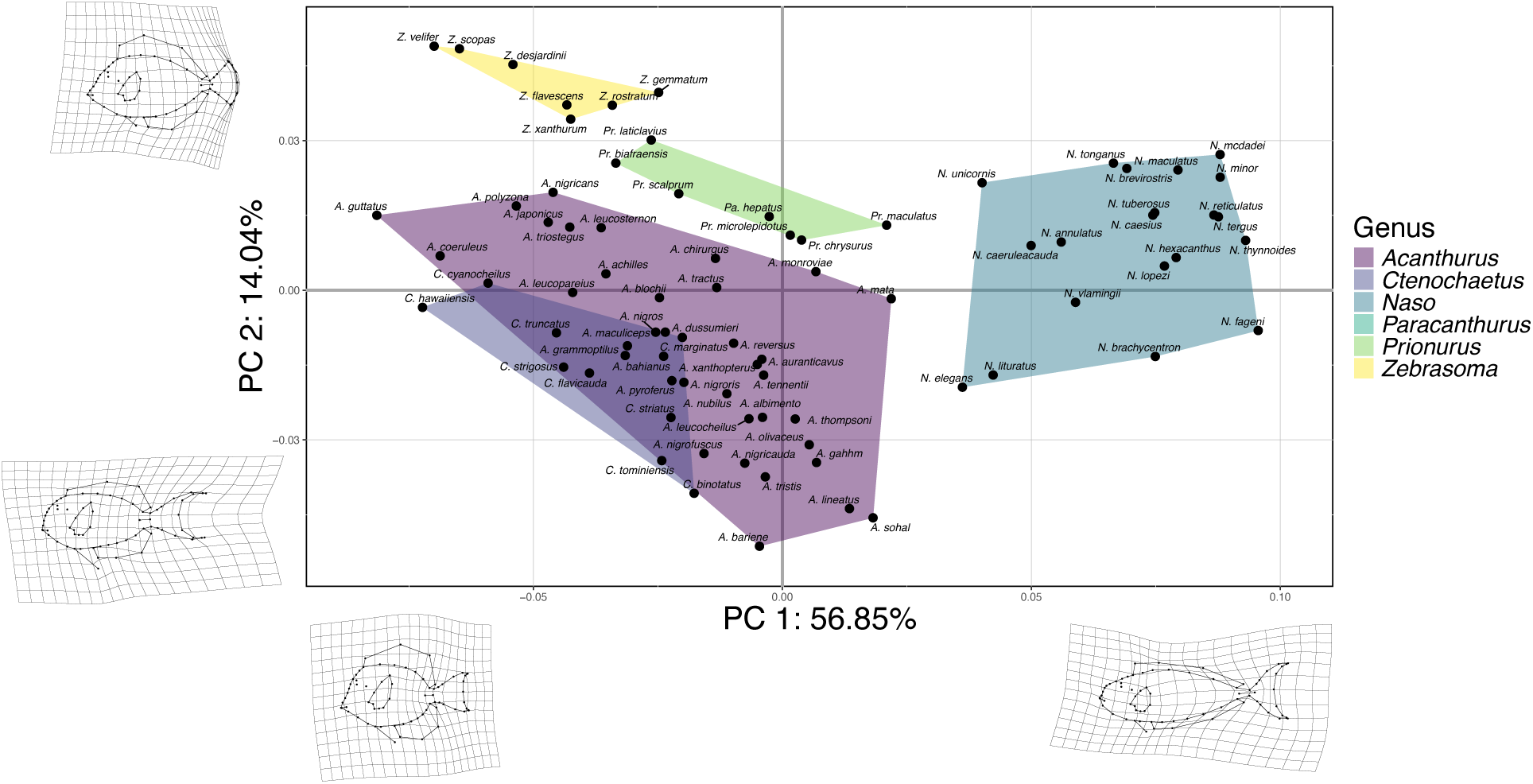
Full Shape Morphospace. PCA plot showing PC1 (56.85%) vs PC2 (14.04%) of the 80 shapes representing each species in the dataset. Genera are indicated by convex hulls. Thin plate splines of the minimum and maximum values of each PC are shown at the corresponding ends of each axis as a deformation of the mean shape of all 80 specimens.

### Phylomorphospace patterns

The first two PCs account for 70.85% of the total variation within our pectoral fin shape dataset, with over 90% being accounted for in the first five PCs. Negative PC1 (45.67%) scores correspond to elongated, high AR pectoral fins and positive scores of PC1 correspond to stout, low AR pectoral fins (Figures 5 & S6). PC2 (25.17%) visualizes a shift from taxa with round pectoral fins with short trailing edges (negative scores) to taxa with sharp pectoral fins with longer trailing edges. *Naso* species exclusively occupy positive PC1 scores corresponding to low pectoral fin ARs (Figure S6). Variation in caudal fin AR seems to be strongly associated with shape variation along PC1 (Figures 5 & S6).

**Figure 5.**
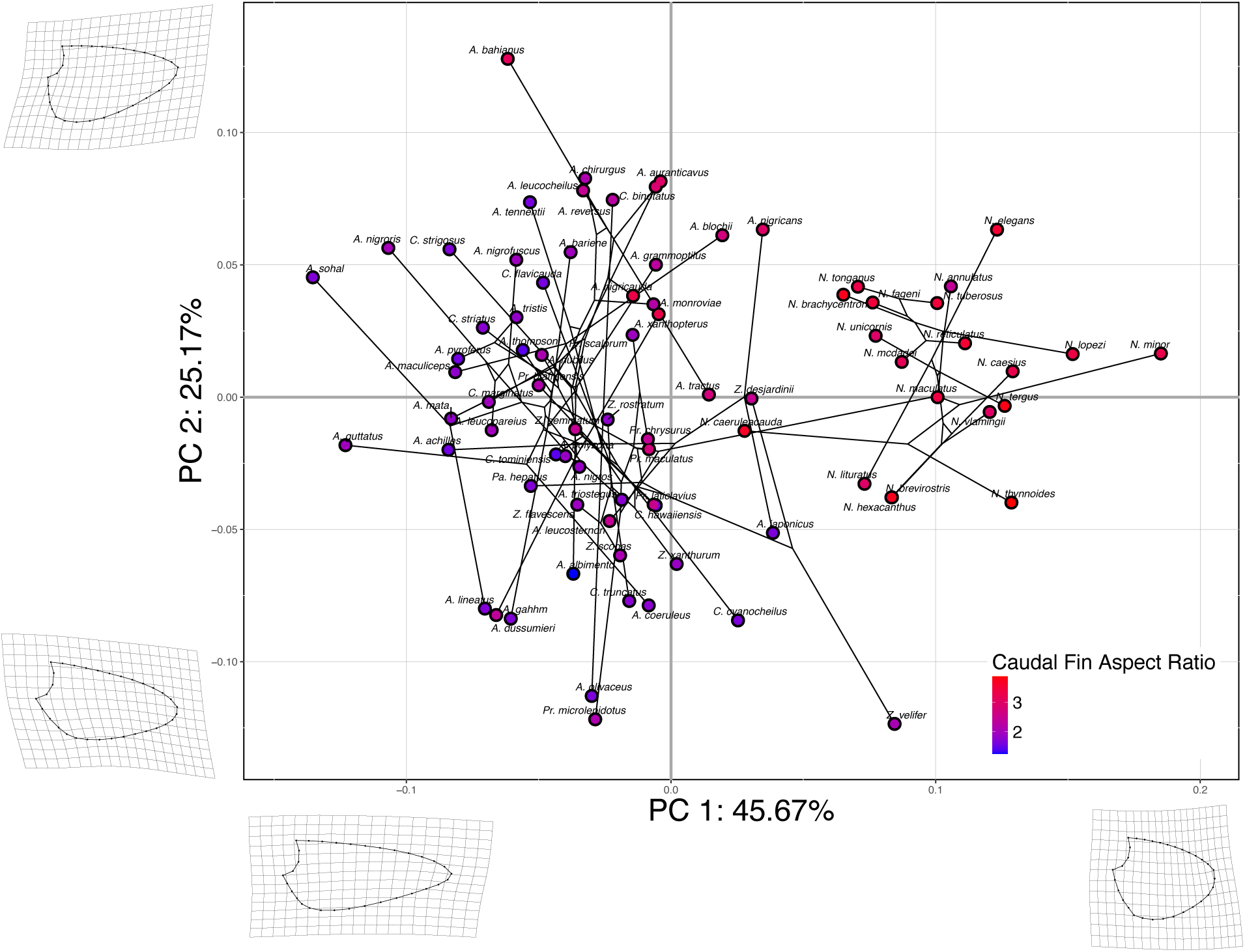
Pectoral Fin Phylomorphospace. PCA plot showing PC1 (45.67%) vs PC2 (25.17%) of the 80 shapes representing the pectoral fin shape of each species in the dataset. Points are color-coded on a gradient of caudal fin AR to visualize potential covariation between caudal fin and pectoral fin shape. Cooler colors represent lower values of caudal fin AR, whereas warmer colors represent higher values. Thin plate splines of the minimum and maximum values of each PC are shown at the corresponding ends of each axis as a deformation of the mean shape of all 80 specimens.

For our caudal fin phylomorphospace, over 90% of the variation in our data is captured in the first four PCs, with the first two containing 86.16% of the overall variation. PC1 (61.15%) describes a shift from concave caudal fins with long trailing ends, curved dorsal and ventral sides, and a wide base (negative scores), to convex caudal fins without exaggerated trailing ends, straight dorsal and ventral sides, and a narrow base (positive scores) (Figures 6 & S7). Negative scores on PC2 (23.01%) correspond to slightly concave caudal fins with a wide base, but no sharp, exaggerated trailing ends, while positive scores correspond to very concave caudal fins with long trailing ends, sharp fin tips, and a narrower base. Variation in pectoral fin AR seems to be more strongly associated with shape variation along PC2, though some variation is also associated with changes along PC1.

**Figure 6.**
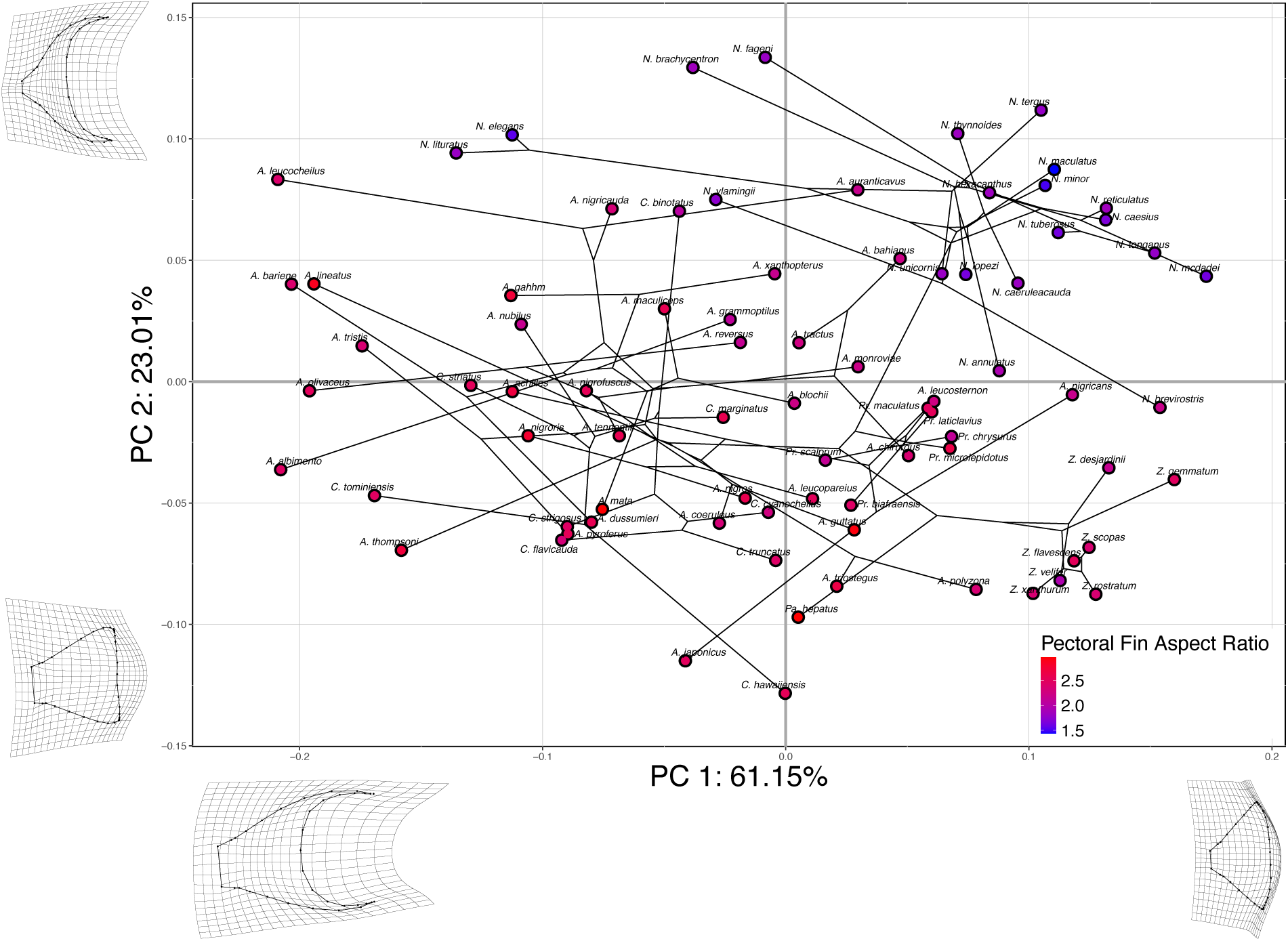
Caudal Fin Phylomorphospace. PCA plot showing PC1 (61.15%) vs PC2 (23.01%) of 79 shapes representing the caudal fin shape of each species in the dataset, with *A. sohal* removed as it is an outlier. Points are color-coded based on a gradient of pectoral fin AR to visualize potential covariation between caudal fin and pectoral fin shape. Cooler colors represent lower values of pectoral fin AR whereas warmer colors represent higher values. Thin plate splines of the minimum and maximum values of each PC are shown at the corresponding ends of each axis as a deformation of the mean shape of all 79 specimens.

The first two PCs of the head shape phylomorphospace make up 57.19% of the overall variation in our dataset with 90% of the variation captured in the first eight PCs. Negative scores of PC1 (40.36%) correspond to longitudinally compressed, but sagittally expanded heads with straight/concave profiles and a pointed snout while positive PC1 scores correspond to longitudinally expanded, but sagittally compressed heads with convex profiles and a rounded snout (Figures 7 & S8). PC2 (16.83%) corresponds to a shift in taxa with posteriorly placed eyes, exaggerated, convex foreheads, and a large mouth (negative scores) to taxa with anterior placed eyes, straight foreheads, and small mouths (positive scores). Dietary ecotype is visually associated with shape variation along PC1 with benthic herbivores/detritivores found in negative PC1 space and omnivores and pelagic planktivores found in positive PC1 space. Three divergent *Acanthurus* species (*A. thompsoni*, *A. mata*, and *A. nubilus*, towards + PC1) are all either omnivorous or pelagic planktivorous unlike the majority benthic herbivorous/detritivorous *Acanthurus* species, and the three divergent *Naso* species (*N. elegans*, *N. literatus*, and *N. unicornis*, towards-PC1) are all benthic herbivorous/detritivorous, unlike the majority omnivorous or pelagic planktivorous *Naso* species.

**Figure 7.**
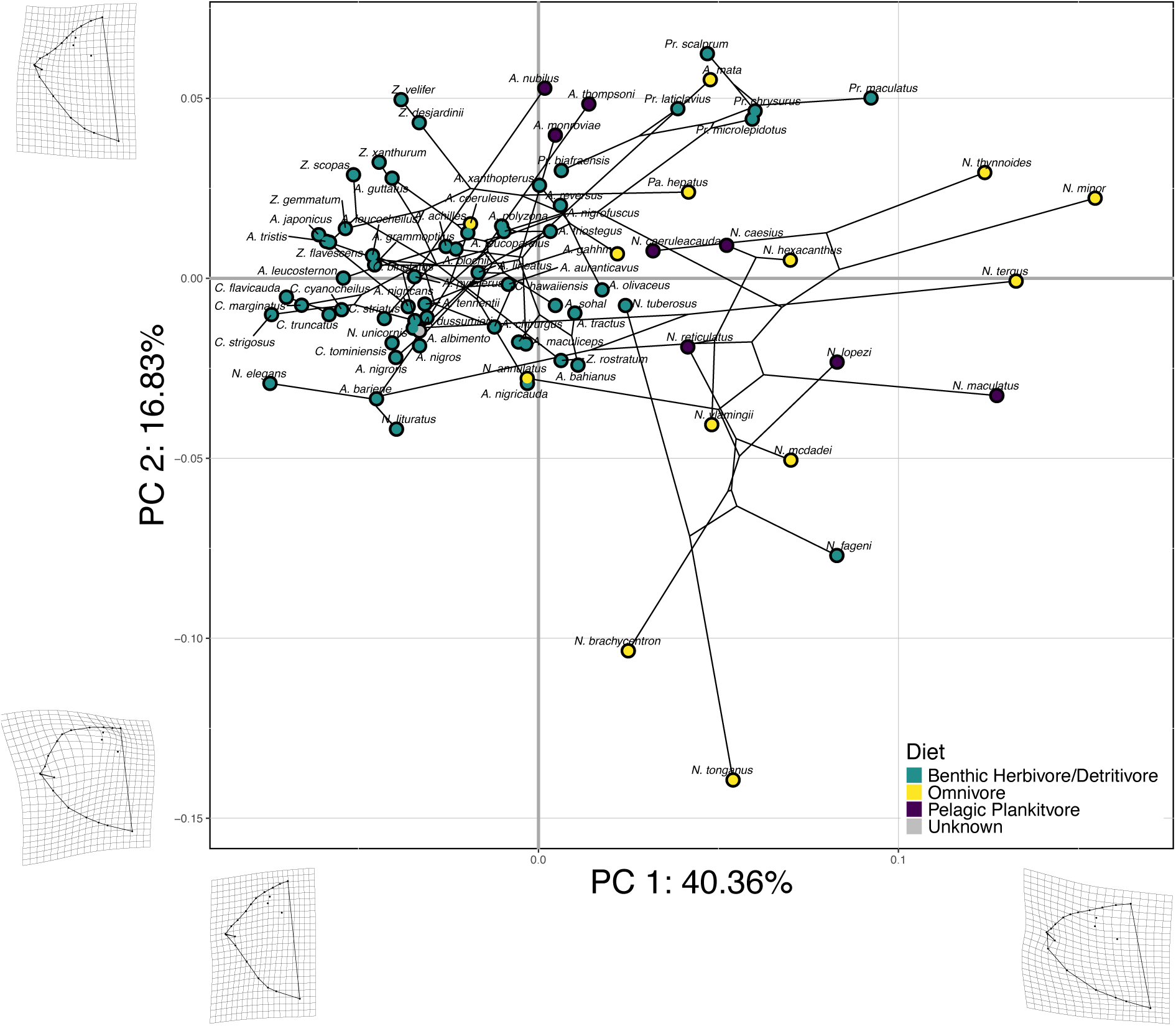
Head Shape Phylomorphospace. PCA plot showing PC1 (40.36%) vs PC2 (16.83%) of 79 shapes representing the head shape of each species in the dataset, with *N. brevirostris* removed as it is an outlier. Points are color-coded by dietary ecotype. Thin plate splines of the minimum and maximum values of each PC are shown at the corresponding ends of each axis as a deformation of the mean shape of all 79 specimens.

Over 90% of the variation in the body shape dataset is contained in the first five PCs, with 81.45% of the overall variation captured by PC1 and PC2. PC1 (74.63%) corresponds to a shift from stout, rounded bodies with wide caudal peduncles and shorter caudal spine areas (negative scores) to elongated bodies with narrow caudal peduncles and longer caudal spine areas (positive scores) (Figure 8). Positive PC2 (6.82%) scores represent square, straight head profiles and shorter caudal spine areas while negative PC2 scores represents convex, round head profiles and longer caudal spine areas. Dietary ecotype is visually associated with shape variation along PC1, but not as strong as seen in our head shape results. While benthic herbivores/detritivores are mostly found in negative PC1 space, the five benthic herbivores/detritivores *Naso* species are found with the other *Naso* species (omnivores and pelagic planktivores, toward + PC1).

**Figure 8.**
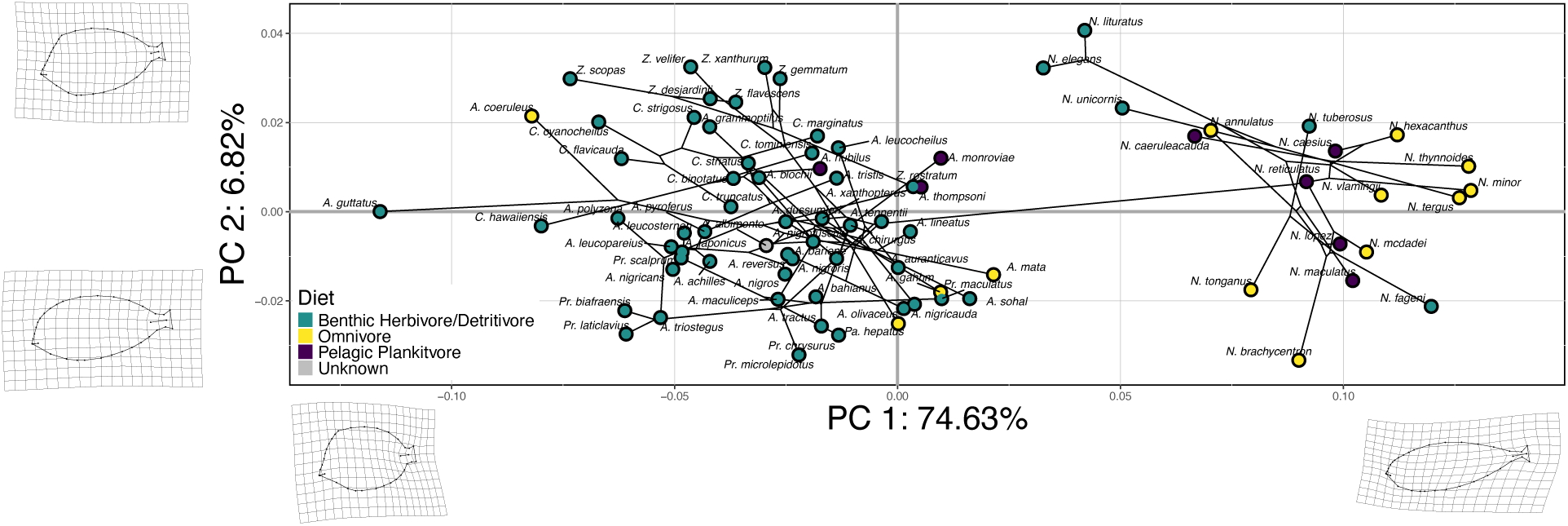
Body Shape Phylomorphospace. PCA plot showing PC1 (74.63%) vs PC2 (6.82%) of 79 shapes representing the body shape of each species present in the dataset, with *N. brevirostris* removed as it is an outlier. Points are color-coded by dietary ecotype. Thin plate splines of the minimum and maximum values of each PC are shown at the corresponding ends of each axis as a deformation of the mean shape of all 79 specimens.

### Ecomorphological relationships

A significant association between both head and body shape variation with dietary ecotype was found, but no significant associations were found between subsequent pairwise comparisons (Supplementary Appendix E), nor between dietary ecotype and our fin shape datasets (Table 1, Supplementary Appendix E). Similarly, a significant association exists between dietary ecotype and elongation ratio (Table 2, Supplementary Appendix E) but not between dietary ecotype and pectoral fin or caudal fin AR. When looking at pairwise tests, benthic herbivores/detritivores significantly differed from both omnivores (p < 0.01) and pelagic planktivores (p < 0.01) in elongation ratio while pelagic planktivores and omnivores did not significantly differ.

**Table 1.** Phylogenetic ANOVA Results for Shape. Summary of the results from each phylogenetic ANOVA testing whether morphological subsets are related to dietary ecotype. The sums of squares (SS), mean squares (MS), R-squared value, F statistic, Z statistic, and p-value are reported for each term. For each term, df = 2. Statistics for the model residuals and the totals can be found in Supplementary Appendix E. Statistically significant relationships between shape and dietary ecotype were based on 1000 iterations for significance testing. (***p < 0.001, **p < 0.01)

|  | SS | MS | R<sup>2</sup> | F | Z | p-value |
| --- | --- | --- | --- | --- | --- | --- |
| **Pectoral Fin Shape** | 0.0002882 | 0.00014411 | 0.02213 | 0.86 | -0.10973 | 0.537 |
| **Caudal Fin Shape** | 0.0003266 | 1.63E-04 | 0.01568 | 0.5975 | -0.61194 | 0.717 |
| **Head Shape** | 0.0011833 | 5.92E-04 | 0.12971 | 5.5892 | 4.0971 | 0.001 *** |
| **Body Shape** | 0.0003852 | 1.93E-04 | 0.08812 | 3.6238 | 2.6753 | 0.005 ** |

**Table 2.** Phylogenetic ANOVA Results for Continuous Metrics. Summary of the results from each phylogenetic ANOVA testing whether continuous fin and body metrics are related to dietary ecotype. The sums of squares (SS), mean squares (MS), F statistic, and p-value are reported for each term. For each term, df = 2. Statistics for the model residuals can be found in Supplementary Appendix E. Statistically significant relationships between shape and dietary ecotype were based on 1000 iterations for significance testing. (***p < 0.001)

|  | SS | MS | F | p-value |
| --- | --- | --- | --- | --- |
| **Pectoral Fin AR** | 1.845254 | 0.922627 | 7.722042 | 0.11 |
| **Caudal Fin AR** | 7.176361 | 3.588181 | 8.811704 | 0.068 |
| **Elongation Ratio** | 5.141448 | 2.570724 | 37.359252 | 0.001 *** |

### Evolutionary dispersion and morphometric correlations

At the extremes, PLS axes exhibit somewhat identical evolutionary variances (pectoral fin:head shape σ ratio = 1.57) to evolutionary variances four times greater in one axis than the other (caudal fin:head shape σ ratio = 4.06). The ratios of evolutionary variances along the primary axes of integration for the significant correlations of caudal fin shape and pectoral fin shape as well as caudal fin shape and body shape were 2.29 and 1.69, respectively. With the Pagel’s λ adjustment, PLS correlation coefficients ranged from 0.34 (pectoral fin shape vs body shape) to 0.59 (caudal fin shape vs body shape), suggesting weak to moderately strong evolutionary correlations between shapes (Table 3, Supplementary Appendix E). Only two of these associations were found to be significant after correcting for multiple comparisons: caudal fin shape vs body shape (Z = 3.67, uncorrected p-value < 0.001, corrected p-value < 0.005, r-PLS = 0.59, Figure 9) and caudal fin shape vs pectoral fin shape (Z = 3.43, uncorrected p-value < 0.001, corrected p-value < 0.005, r-PLS = 0.52, Figure 10). The strength of integration between these two tests did not differ significantly (Supplementary Appendix E). No other tested shape pairs exceeded the significance level with the Holm-Bonferroni correction. Importantly, all comparisons were significant, and the levels of correlation increased under the “all or nothing” assumptions compared to our Pagel’s λ adjustment results (Table S4).

**Figure 9.**
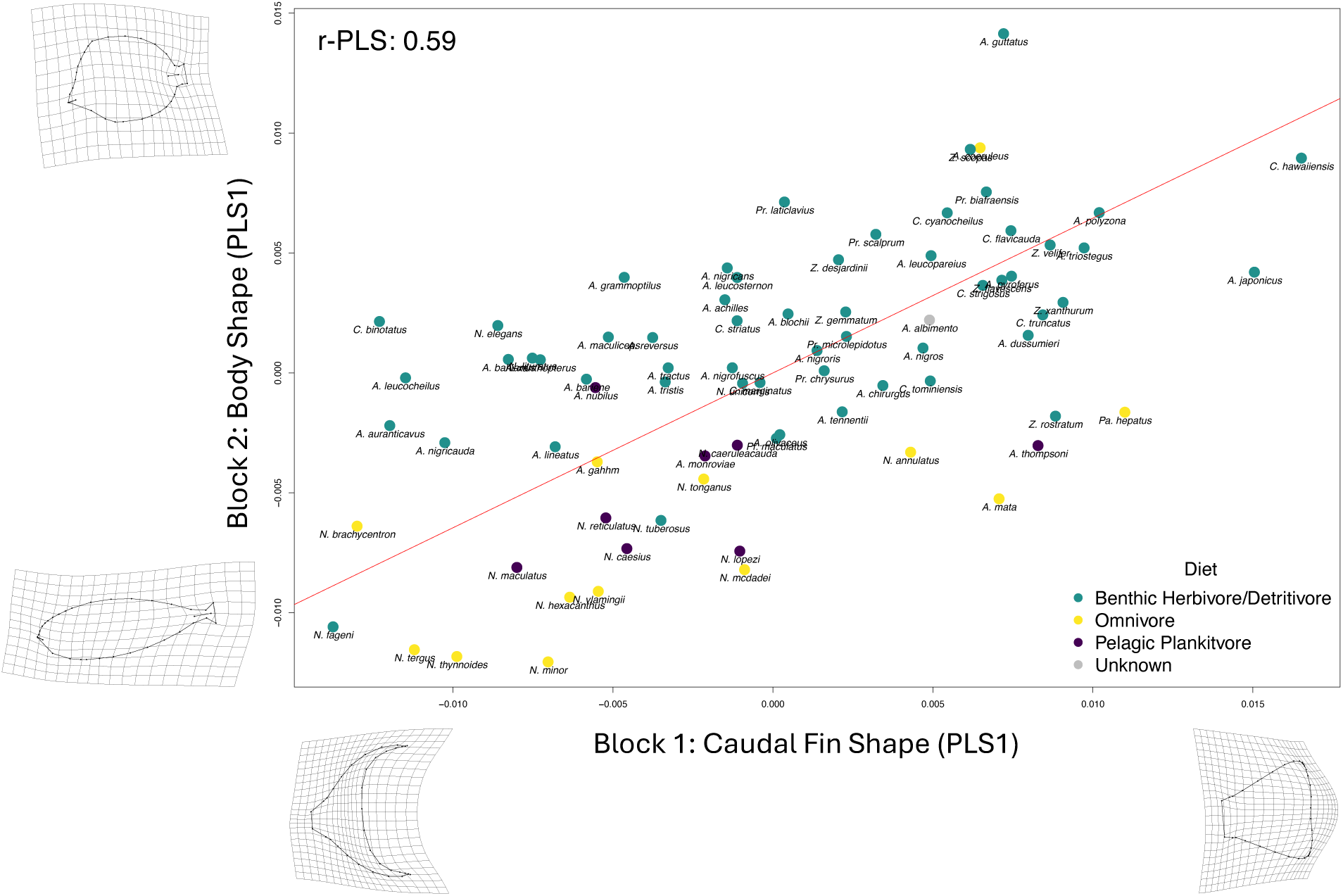
Caudal Fin Shape vs Body Shape 2B-PLS. Plot showing the first PLS scores for caudal fin shape (block 1) and body shape (block 2). Caudal fin shape and body shape are significantly paired (p-value < 0.01), with a PLS correlation coefficient of 0.59. The red line indicates the correlation between the two PLS axes yielded by a phylogenetically-informed PLS test. Thin plate splines at the axes’ extremes visualize shape change across their respective axes. Points are color-coded based on dietary ecotype.

**Figure 10.**
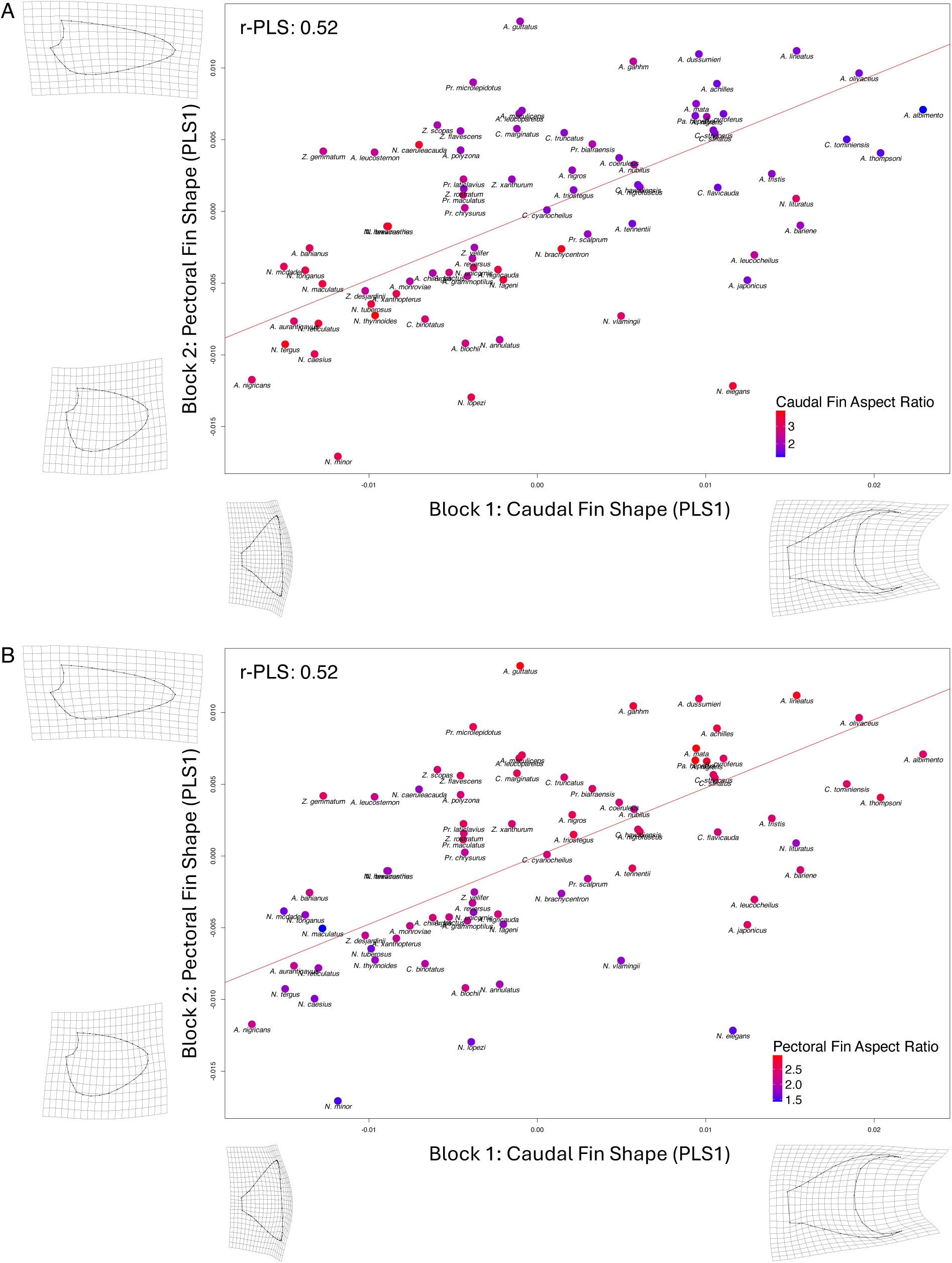
Caudal Fin Shape vs Pectoral Fin Shape 2B-PLS. Plots showing the first PLS scores for caudal fin shape (block 1) and pectoral fin shape (block 2). Caudal fin shape and pectoral fin shape are significantly paired (p-value < 0.01), with a PLS correlation coefficient of 0.52, indicating a moderately strong correlation between shapes. Pectoral fin and caudal fin shape are inversely related: low/high AR pectoral fins are correlated with high/low AR caudal fins. The red line indicates the correlation between the two PLS axes yielded by a phylogenetically-informed PLS test. Thin plate splines at the axes’ extremes visualize shape change across their respective axes. Points are color-coded on a gradient to indicate mean (A) caudal fin AR and (B) pectoral fin AR for each species. Cooler colors represent lower values of fin ARs, whereas warmer colors represent higher values of fin ARs.

**Table 3.** 2B-PLS Results. Summary of the results from 2B-PLS tests with our Pagel’s λ adjustment done on each morphological subset pairing (except head shape vs body shape). Raw p-values and Holm-Bonferroni-corrected p-values are stated, along with the PLS correlation coefficients (r-PLS) and effect sizes (Z) for each comparison. The λ value that maximized the likelihood of the data (PLS scores) used to rescale the phylogeny is reported. Corresponding log likelihood and AIC scores can be found in Supplementary Appendix E. The evolutionary variances along each PLS axes are noted (PLS X1 σ and PLS Y1 σ, respectively) as well as the evolutionary covariation between the two PLS axes and the ratio of evolutionary variances. (**p < 0.01, *p < 0.05)

| Block 1 Shape | Block 2 Shape | p-value (raw) | p-value (corrected) | r-PLS | Effect Size (Z) | $\lambda$ | PLS X1 $\sigma$ | PLS Y1 $\sigma$ | Cov(PLS1) | $\sigma$ ratio |
| --- | --- | --- | --- | --- | --- | --- | --- | --- | --- | --- |
| Caudal | Pectoral | 0.001** | 0.005** | 0.522 | 3.4304 | 0.2 | 9.43E-05 | 4.13E-05 | 3.26E-05 | 2.285 |
| Caudal | Head | 0.027* | 0.081 | 0.431 | 1.9684 | 0.15 | 1.25E-04 | 3.08E-05 | 2.67E-05 | 4.064 |
| Pectoral | Head | 0.051 | 0.102 | 0.446 | 1.6347 | 0.19 | 4.58E-05 | 2.92E-05 | 1.63E-05 | 1.567 |
| Caudal | Body | 0.001** | 0.005** | 0.588 | 3.6737 | 0.17 | 4.79E-05 | 2.83E-05 | 2.16E-05 | 1.691 |
| Pectoral | Body | 0.194 | 0.194 | 0.335 | 0.903 | 0.45 | 3.9E-05 | 2.18E-05 | 9.76E-06 | 1.793 |

## Discussion

We aimed to investigate the ecological associations and evolutionary covariations within surgeonfishes by revising phylogenetic relationships, describing morphological patterns, and presenting a solution to a current problem in phylogenetic comparative methods. We found that surgeonfish head and body shapes are significantly related to their dietary ecotype, and they have evolved an inverse relationship between caudal fin and pectoral fin AR. Furthermore, shared evolutionary history does not fully explain the significant shape correlations seen in surgeonfishes, which all appear to relate to the caudal fin. Below, we detail our insights into surgeonfish ecomorphology and shape evolution using new tools for morphometrics and comparative methods and discuss the need for more investigation into the relationship between differences in the evolutionary variance along the primary axes of integration and degree of covariation across shapes.

### Phylogenetic relationships and the ecomorphology of head and body shape

Our molecular phylogenetic analysis of surgeonfishes is the most complete in terms of taxonomic sampling and number of loci, encompassing 97% of surgeonfish diversity and adding 10 additional loci. Our new molecular phylogeny confirms many of the relationships previously proposed in several morphological trees (Guiasu & Winterbottom, 1993; Winterbottom, 1993) and expands the conclusions made in previous molecular phylogenies (Clements et al., 2003; Klanten et al., 2004; Sorenson et al., 2013, details in Supplementary Discussion).

Results from the phylomorphospaces and statistical tests show that head and body shape are significantly associated with fishes’ dietary ecotype. Benthic herbivores/detritivores tend to have more elongated bodies, small mouths, straight foreheads, and short heads (Figures 7, 8, & S8) while omnivores and pelagic planktivores both exhibit deeper bodies, larger mouths, humped foreheads, and longer heads (Figures 7, 8, & S8). Head shape and body elongation are functionally important traits of fishes as they are associated with habitat, trophic niches, and feeding modes, potentially limiting the range of possible head and body morphologies fishes can exhibit and affecting evolutionary rates (Bejarano et al., 2017; Claverie & Wainwright, 2014; Cooper & Westneat, 2009; Floeter et al., 2018; Friedman et al., 2016, 2019; López-Fernández et al., 2013; Mahe et al., 2014; Siqueira et al., 2020; Tavera et al., 2018). Specifically in surgeonfishes, body and head shape (Robertson et al., 1979) and related mechanics (Mihalitsis & Wainwright, 2024; Perevolotsky et al., 2020) have been hypothesized to facilitate some species’ specialized herbivorous feeding mode. The smaller mouths and more maneuverable body of benthic herbivores/detritivores may be advantageous in their distinct grazing behavior, while the larger mouths of pelagic planktivores may maximize dietary intake. This relationship has been discussed in a larger reef fish study (Claverie & Wainwright, 2014). We see that omnivores do not exhibit an “intermediate” head and body shape despite these taxa occupying a broader trophic niche. One explanation for this result could be that an intermediate shape (say a mid-sized mouth and a body not optimized for either maneuverability or steady swimming) does not perform well with specific behaviors related to benthic herbivorous/detritivorous feeding or pelagic planktivory, making it an unstable morphology in terms of evolutionary fitness. Why then do omnivores tend to exhibit the pelagic planktivore phenotype over the benthic herbivore/detritivore phenotype? Perhaps, pelagic planktivore dietary ecotype is more specialized; fishes can be successful benthic herbivores/detritivores with a larger mouth and elongated bodies but would be unsuccessful at pelagic planktivory with smaller mouths and bodies not optimized for steady swimming performance (Langerhans, 2008). However, our study found no significant pairwise comparisons between dietary ecotypes; more investigation is needed to understand the exact morphological specializations related to dietary ecotype in this group.

While dietary ecotype may impact head and body shape, it has relatively weak explanatory power (Table 1). Other factors could be contributing to body and head shape variation in surgeonfishes. For example, the patterns described in the previous paragraph could be driven by clade-specific morphologies. *Naso*, which consists mostly of omnivores and pelagic planktivores, cluster in body shape morphospace regardless of dietary ecotype. However, clade relationships may not impact head shape evolution as much, as most of the benthic herbivorous/detritivorous *Naso* species do not cluster with omnivorous and pelagic planktivorous *Naso* species. Future ancestral state estimation studies could reveal that these omnivores have evolved from pelagic planktivore lineages and are retaining an ancestral shape.

### Evolution of inverse fin shape related to locomotion

The main axis of variation in pectoral fin morphology across many fishes is the distinction between high AR (> 2.0, Walker & Westneat, 2000; Westneat et al., 2017), flapping fins that facilitate efficiency and speed during swimming, versus low AR, paddling fins that are specialized for maneuverability (Gerstner, 1999; Walker & Westneat, 2000, 2002). Our fin shape analyses show that *Acanthurus*, *Ctenochaetus*, *Paracanthurus*, *Prionurus*, and *Zebrasoma* tend to have high AR, wing-like pectoral fins well suited for flapping kinematics. In contrast, members of the genus *Naso* have low AR, paddle-shaped pectoral fins that maneuver effectively, a potential advantage for their pelagic lifestyle. The key biomechanical paradigm of thrust efficiency vs maneuverability in pectoral fin shape found in wrasses and parrotfishes is operating in surgeonfishes as well, yet few biomechanical analyses of pectoral fin locomotion have been performed in surgeonfishes (Mihalitsis & Wainwright, 2024; Perevolotsky et al., 2020; Schakmann & Korsmeyer, 2023). Future work on pectoral kinematics integrated with musculoskeletal design and neural control (Drucker, 1996; Flammang & Lauder, 2013; Westneat, 1996) during steady swimming and maneuvering behaviors may yield important insights into locomotor trade-offs in acanthurids.

A similar, but inverse, relationship is found across surgeonfish caudal fin morphology. Caudal fins range from high AR with narrow caudal peduncles, which are exhibited by many *Naso* species, to low AR with wider peduncles, which are more common in other genera. High AR caudal fins have been shown to increase the lift-drag ratio during swimming, allowing for efficiency and greater endurance swimming speeds (Sambilay, 1990), while low AR fins are associated with high-speed sprint swimming (Webb, 1982; Weihs, 1973). Narrower peduncles and some degree of forking (as opposed to no forking, or extreme forking) may also allow for enhanced propulsive performance (Krishnadas et al., 2018). Other research has indicated that crescent-shaped caudal fins (another high AR shape) yield less thrust than their less-curved counterparts but are more efficient when cruising (Chang et al., 2012). *Naso* species rely on their caudal fin, rather than pectoral fin, for propulsion. Thus, their caudal fins may be specialized for enhanced and efficient locomotion while cruising in an open pelagic environment (Klanten et al., 2004, Supplementary Appendix C). Exploration of swimming kinematics and red muscle/axial tendon morphology (Westneat & Wainwright, 2001) on high-performance axially swimming surgeonfishes may provide a holistic view of biomechanical specializations.

Combining our fin shape results, we see that most surgeonfishes have evolved two alternative combinations of propulsive fin shape, one with a high AR caudal fin and low AR pectoral fins, and the other with high AR pectoral fins and a low AR caudal fin (Figures 3, 5, 6), although it is intriguing to note that many species show intermediate ARs and are scattered along the continuum (Figure 10). The evolution of a high/low AR tail with low/high AR pectoral fins has biomechanical and locomotor significance, with the former pairing associated with body-caudal fin propulsion and the latter with labriform propulsion. *Acanthurus* species employing labriform propulsion are some of the fastest fishes according to field measurements of typical, steady, swimming behaviors in wave-swept reef areas (Fulton, 2007; Fulton et al., 2005). However, *Naso* and some other species that primarily oscillate the tail for propulsion are also high-performance swimmers (Fulton, 2007; Fulton et al., 2005).

Expanding these ideas broadly across reef fishes could reveal large-scale patterns of caudal and pectoral fin shape correlated evolution. Pectoral fin AR has been measured in 46 species of damselfishes (Pomacentridae) and ranges from 0.63 to 1.67 (Fulton et al., 2005), suggesting these fishes exhibit lower pectoral fin ARs. The family Labridae, with over 600 species, is the only other reef fish family for which pectoral AR measures are fairly comprehensive (Aiello et al., 2017; Wainwright et al., 2002; Westneat et al., 2017). Labrids have more variation in their pectoral fin ARs than surgeonfishes and damselfishes; values range from 0.8 to 4.6 across 340 species, with over 150 species exhibiting high AR wing-like fins. Investigating caudal fin AR and fin shapes across these taxa and including groups more closely related to acanthurids like the angelfishes and butterflyfishes will reveal if inverse fin shape relationships have evolved across reef fishes in general, or if they are a defining characteristic of the acanthurids.

### Evolutionary covariation between shapes

We suggest that our Pagel’s λ adjustment is a more conservative approach to the standard assumptions of shape covariation, due to the differences in resulting significance and correlation values (Table S4). Significant shape correlations between morphometric subsets within Acanthuridae indicate that the evolution of one shape may influence the evolution of another in a complementary pattern. While our Pagel’s λ adjustment results support strong correlations between the caudal and pectoral fins as well as caudal fin shape and body shape, the shared evolutionary histories of species have low explanatory power towards the patterns of covariation we found between shapes. Other factors, such as dietary ecotype or swimming biomechanics of surgeonfishes, could be driving these correlations.

The correlation between caudal fin and body shape (Figure 9) shows that the omnivores and pelagic planktivores separate from benthic herbivores/detritivores in PLS1 space, mostly along the axis relating to body shape, which aligns with our body shape results discussed above. However, some omnivorous species seem to separate from pelagic planktivorous species along both axes, suggesting that combinations of certain body and caudal fin morphologies may distinguish all three dietary ecotypes from one another in surgeonfishes. Morphological integration (Burns et al., 2023; Evans et al., 2021; Larouche et al., 2018) and dietary ecotype traits (Friedman et al., 2016; McCord et al., 2021; Ribeiro et al., 2018, Santaquiteria et al., 2026) have been separately linked to patterns of diversification, rates of evolution, and adaptive evolutionary models. Thus, dietary ecotype has unexplored impacts on the evolution of shape covariations. In surgeonfishes, dietary ecotype diversity could promote covariation in shapes but ultimately result in similar individual shapes across certain dietary ecotypes, especially related to the caudal fin.

Dietary ecotype could be contributing to the covariation between caudal fin and body shape due to functional demands related to ecological feeding behavior, but more work is needed to confirm this. Coordinated movements across fins and bodies facilitate feeding behavior in wrasses and parrotfishes (Rice et al., 2008; Rice & Westneat, 2005), and studies have shown that herbivorous surgeonfishes have unique feeding behaviors that involve coupled movements of the head, body, and fins (Mihalitsis & Wainwright, 2024; Perevolotsky et al., 2020). Considering fin and body shape enable different locomotor modes in fishes (Friedman et al., 2021; Gerstner, 1999; Walker et al., 2013; Walker & Westneat, 2000, 2002), our results present evidence that the observed correlation in caudal fin shape and body shape may have arisen to meet the locomotor/functional demands related to specific dietary ecotypes. Stayton (2019)’s adaptive landscape study showed that species distribution in morphospace overlaps with the distribution in performance optima. Integrated systems also score higher in performance landscapes than non-integrated systems, and diet is suggested to be the mechanism of this covariation (Klingenberg, 2013; Sansalone et al., 2022). However, other factors could be driving the covariation in surgeonfishes. Much of the separation in PLS space between dietary ecotypes is related to body shape, which, as previously discussed, may be constrained due to evolutionary ancestry. Figure 9 shows that *Naso* seems to cluster in PLS space, with some exceptions, and most of the omnivorous species that separate from the pelagic planktivores species are not *Naso* species. Future studies investigating the impact of *Naso* on the covariation between body and caudal fin shape is needed.

Looking at our PLS results for the covariation between the caudal and pectoral fin (Figure 10), we provide further evidence for the evolution of pectoral fin and caudal fin shapes as pairs of low/high AR (Figures 3, 5, 6). This correlation between pectoral and caudal fin shapes does not seem to be associated with dietary ecotype (Figure S9); likely, evolutionary covariation among these propulsive fins is driven by the biomechanical demands of different swimming modes (labriform or body-caudal-fin). However, surgeonfishes exhibited extreme, inverse pairings of high/low AR fins as well as pairings of intermediate AR fins, suggesting that a wide range of fin shapes and locomotor mechanisms are retained in the family and contribute to high morphological diversity in surgeonfishes.

Morphological integration arising from functional coupling between shapes is expected to promote evolutionary correlations (Cheverud, 1982, 1984; Olson & Miller, 1958; Riedl, 1978; Schlosser & Wagner, 2004). Classic thoughts on shape covariation suggest integrated shapes may be evolutionarily restrictive and prevent rapid diversification, while non-integrated shapes can evolve independently and may be more likely to promote rapid diversification (Wagner & Altenberg, 1996), although this depends on the lability of the structure of integration (Webster & Zelditch, 2011). However, recent studies have suggested that integration may be related to rapid diversification, especially in fishes (Burns et al., 2023; Evans et al., 2021; Larouche et al., 2018). Shape covariation has implications for overall diversification rates, but what evolutionary variance patterns exist *between* compared shapes? And do the observed relationships depend on the strength of shape correlation? In all of our 2B-PLS tests, evolutionary variances along the primary axes of integration differed across shapes by at least a factor of 1.5 (Table 3). In our significantly correlated shapes, PLS scores related to caudal fin shape exhibited evolutionary variances 1.7 to 2.3 times that of PLS scores related to body and pectoral fin shape, respectively, suggesting that even when morphological shapes are correlated, their evolutionary variances along the primary axis of integration can differ greatly.

However, future work might explore whether evolutionary variances between shapes along the primary axis of integration are related to the strength of their correlation, as no consistent pattern was found in this study. For example, while our shapes with the most similar evolutionary variances along the primary axis of integration did not significantly covary with one another (Table 3), they were more correlated than the shapes that exhibited the largest difference in evolutionary variances, which also did not significantly covary with one another. Additionally, while the strongly correlated caudal fin and pectoral fin shapes exhibited the second largest difference in evolutionary dispersion along the primary dimension of integration, the strongest association of caudal fin shape and body shape exhibited the second lowest difference in evolutionary dispersion. Our novel methodology, which is a more conservative approach than the standard assumptions of shape covariation, can be used in future work exploring the relationship between evolutionary morphometric variation along the primary axis of integration and the strength of their related shape associations, especially in functionally and ecologically important systems that are not evolving under BM. It is important to note that we aren’t comparing evolutionary rates between blocks, because our PLS scores are data-derived latent variables which are inherently statistical and not biological. Thus, while evolutionary rate differences *could* be causing the patterns we see, many other processes could be contributing as well, such as rate variation between blocks, different constraints, or variation in the strength of selection. Future studies investigating the contribution of these phenomena to the patterns we present should be explored.

## Conclusion

Here, we offer a solution to a long-standing problem in phylogenetic comparative methods through a Pagel’s λ adjustment to estimate the influence of shared evolutionary history on shape covariation within the surgeonfishes (Acanthuridae) and uncover important ecomorphological patterns and correlations. Surgeonfishes head and body shape diversity are associated with dietary ecotype, and locomotor demands likely contribute to surgeonfishes exhibiting alternative pairs of high/low aspect ratio (AR) tails with low/high AR pectoral fins, respectively. While caudal fin shape covaries with both body and pectoral fin shape, phylogenetic signal is generally low, indicating that the shared evolutionary history of species does not substantially explain these major covariations between shapes among surgeonfishes. This covariation may be related to their ecology and behavior; caudal fin and body shape covary and relate to dietary ecotype, whereas the covariation between the caudal fin and pectoral fin shape relates to locomotor mechanics. The caudal fin shape exhibits the highest degree of evolutionary variance along the primary axis of integration and is strongly associated with almost all other shape subsets, demonstrating the importance of investigating the relationship between the evolutionary dispersion along the primary dimension of integration of morphometric evolution and the strength of association between morphometric shapes. Finally, without estimating the influence of phylogeny on shape covariances, studies may not adequately estimate evolutionary rate matrices and transform their data to remove phylogenetic signal, potentially resulting in biased estimates of shape correlation. When we limited our analyses to the assumption that shape covariances between taxa are either independent of (λ=0), or completely described by (λ=1), their shared evolutionary history, each shape correlation increased and was significant, suggesting that both the “all” or “nothing” influence of phylogeny on trait covariances are less conservative assumptions. Our novel approach may provide future researchers a broader array of phylogenetic tools to help avoid bias by estimating the influence of shared evolutionary history on trait covariation.

## Supplementary Material

Supplementary material is available online at *Evolution*.

## Data Availability

The data underlying this article are available in figshare with the following link:https://figshare.com/projects/Phylogenetic_Relationships_and_the_Evolution_of_Fin_and_Body_Shape_in_the_Surgeonfishes_-_Supplementary_Materials/259151. SurgeonShape can be downloaded at the author’s GitHub using the following link: https://github.com/mwestneat.

## Authorship Contribution

**Linnea L. Lungstrom**: Conceptualization, Data curation, Formal analysis, Investigation, Methodology, Visualization, Writing – original draft. **Mireille Farjo**: Conceptualization, Data curation, Investigation, Writing – review & editing. **Ryan Isdonas**: Data curation, Investigation, Writing – review & editing. **Andrew B. George**: Data curation, Investigation, Writing – review & editing. **Mark W. Westneat**: Conceptualization, Data curation, Formal analysis, Funding acquisition, Methodology, Software, Supervision, Visualization, Writing – review & editing.

## Acknowledgements

Thank you to the staff of the Division of Fishes at the Field Museum (FMNH) (Caleb McMahan, Susan Mochel, and Kevin Swagel) for providing access to the collections for specimen imaging. We are also grateful to many institutional and online databases (Smithsonian National Museum of Natural History, Bishop Museum, FishBase, John E. Randall’s and Jeffrey T. Williams’ Fish Photos) that contributed photos to our dataset. Thank you to Mark Webster for advice on morphometric approaches and Graham Slater for discussions about phylogenetic corrections related to morphometric analysis. We also thank G. Slater as well as Stephen Pruett-Jones and Dylan Hubl for critical comments on the initial manuscript. Special thanks to Lydia Smith, Jillian Henss, and Charlene McCord for gene sequencing work at FMNH.

