## Supplementary for "Phylogenetics, Trait Covariance Analysis, and the Evolution of Fin and Body Shape in the Surgeonfishes"

**Supplementary Methods** (p. 1)

**Supplementary Results** (p. 7)

**Supplementary Discussion** (p. 9)

**Supplemental Figures** (p. 13):
1. SurgeonShape App
2. Maximum-likelihood Tree for Acanthuridae
3. Estimated Ages for All Nodes in New Phylogeny
4. Posterior Probabilities for All Nodes in New Phylogeny
5. Elongation Ratio along Acanthuridae Phylogeny
6. Pectoral Fin Phylomorphospace
7. Caudal Fin Phylomorphospace
8. Head Shape Morphospace
9. Caudal Fin Shape vs Pectoral Fin Shape Two-block PLS

**Supplemental Tables**:
1. Substitution Models (p. 22)
2. Landmarks (p. 23)
3. Curves (p. 24)
4. 2B-PLS Results across Methodologies (p. 25)

**Appendices**:

A. **GenBank Sources.** Sources of genetic data for phylogenetic analysis of the Acanthuridae, with specific contributions of prior studies color coded for clarity of attribution of authorship in the data used in this study. figshare link

B. **Image Sources.** Sources of surgeonfish images used for morphometric analysis of the Acanthuridae. figshare link

C. **Trait References.** Citation sources for ecological traits coded for surgeonfishes. figshare link

D. **Elongation and Fin Aspect Ratio Data.** Raw ratio values for each image and mean ratio values for each surgeonfish species. figshare link

E. **Statistical Report.** Full statistics related to phylogenetic ANOVAs and two-block PLS tests. figshare link

**Files:**
1. Acanth53BEAST.xml – beast XML figshare link
2. Acanth53CombinedTree.tre – MCC tree from combined trees figshare link
3. Coordinates.txt – coordinate data exported from SurgeonShape figshare link
4. Code.R – code used to run read in data, run analyses, build plots, etc. figshare link

### Supplementary Methods

*DNA sequence generation and alignment*

Tissues and/or data for only three extant species of *Acanthurus* were unavailable: *Acanthurus albipectoralis*, *A. chronixis*, and *A. fowleri*. The total count of 83 surgeonfishes used here differs from the FishBase.org list of 84 taxa. We reject *A. randalli* and *Prionurus punctatus*; although they remain in FishBase, they are synonyms of *A. bahianus* (Smith-Vaniz et al., 2002) and *P. laticlavius* (Ludt et al., 2019), respectively. We include both *A. nigroris* and *A. nigros* (Randall et al., 2011); *A. nigros* is widespread throughout the Pacific but is not yet listed by FishBase. Portions of 19 genes were analyzed, including eight mitochondrial regions (12S, 16S, ND3, ND4, ND5, ND6, COI, and CYTB) and 11 nuclear markers (ETS2, RAG1, RAG2, TMO4C4, DLX2, OTX1, BMP4, MYH6, PLAGL2, ZIC1, and RHOD). Many of the data analyzed here were obtained from previous studies (Clements et al., 2003; Fessler & Westneat, 2007; Holcroft & Wiley, 2008; Klanten et al., 2004; Ludt et al., 2015; Near et al., 2013; Sorenson et al., 2013; Appendix A), but we contribute a large set of new DNA sequences for several genes for both acanthurids and outgroups.

New DNA sequences were generated using standard extraction and Sanger sequencing approaches outlined in previous phylogenetic studies (McCord & Westneat, 2016). Briefly, DNA was extracted from muscle or gill tissue, and double-stranded DNA products were amplified using polymerase chain reaction (PCR) from DNA isolates of the 12S, 16S, ND3, TMO4C4, RAG1, RAG2, BMP4, and DLX2 gene fragments. PCR products were then cycle-sequenced and purified. Then, sequences of both strands were generated on an ABI PRISM 3730 Genetic Analyzer (Applied Biosystem, Foster City, CA). Primers used for DNA amplification and sequencing are identical to previous studies (McCord & Westneat, 2016).

Sequences for 12S and 16S were manually aligned to secondary structure models for ribosomal sequences using previously published homologous sequences (Fessler & Westneat, 2007), whereas protein coding genes were automatically aligned using the ClustalW or Muscle modules within Mesquite (Maddison & Maddison, 2025). Some sequences were trimmed to the size of the smallest fragment to minimize the amount of missing data in the data matrix. Gaps and ambiguously aligned regions were excluded from analysis. After trimming sequence ends, the character count for each gene was 12S (925), 16S (575), ND3 (363), ND4 (1622), ND5 (1847), ND6 (520), COI (652), ETS2 (478), CYTB (1093), RAG1 (1443), RAG2 (802), TMO4c4 (514), BMP4 (481), DLX2 (518), OTX1 (671), MYH6 (768), PLAGL2 (761), ZIC1 (821), and RHOD (798) for a total of 15,669 nucleotide characters.

*GenBank errors*

Here, we note several GenBank sequences that were not used due to potential errors in identification or taxonomic changes. Accession KJ679904 listed as COI for *Acanthurus tristis* appears to be a misidentified *A. triostegus*. Accession PQ412526 voucher USNM:AG7PW17 *A. reversus* whole mitochondrion BLASTS to *Naso* species. Data for *Prionurus punctatus* should be synonymized with *P. laticlavius*. GenBank accessions for *A. nigroris* and *A. nigros* can only be disambiguated by having the specimen locality and preferably a voucher specimen, as the more widespread localities are *A. nigros*, and *A. nigroris* is now restricted to the Hawaiian Islands and Johnston Island.

*Phylogenetic analysis and priors*

ModelFinder (IQ-TREE version 2; Kalyaanamoorthy et al., 2017; Minh et al., 2020) was used to identify the best fitting models of nucleotide evolution for gene sequences by starting with the full partition model (19 separate gene partitions) and merging genes into single partitions until the model fit did not increase further (Kalyaanamoorthy et al., 2017). Using this best-fit partition model, IQ-TREE reconstructed the maximum likelihood tree. This maximum likelihood tree was then imported into Mesquite (Maddison & Maddison, 2025) where the tree was ultrametricized. Branch lengths were then scaled so that no branch length was 0 and that nodes with known fossil calibrations were consistent with their known ages. The tree was again ultrametricized and then exported to be used as the starting tree in BEAUti 2.7.7. along with the resulting models of nucleotide evolution for each partition (Table S1) obtained from IQ-TREE.

We selected the optimised relaxed clock (ORC) model in BEAUti, which is based on a log normal distribution of rates which employs an adaptive operator that learns the best weight of other operators and Bactrian proposal kernels that improve parameter space exploration. These features of the model improve speed of convergence and mixing in the Markov Chain Monte Carlo (MCMC) analysis. We used the standard birth death model as a tree prior, with a relative death rate of 0.5 and a birth diff rate of 1.0. A net diversification rate (BDBirthRate prior) and extinction fraction (BDDeathRate prior) were modeled as an exponential (mean = 0.1) and beta (alpha = 2, beta = 1) distribution respectively. We also used two fossil calibrations for the ORC model. The Middle Eocene *Sorbinithurus sorbinii* from Monte Bolca (Italy), which was identified by Tyler (2000) as a stem member of the Nasinae, was used to assign a minimum age of 48.5 Ma (Papazzoni & Trevisani, 2006) to the crown Acanthuridae. For dating the split of acanthurids from luvarids, we used the Eocene fossil *Kushlukia permira* from the Ypresian

(Lower Eocene), identified by Bannikov and Tyler (1995) as a stem luvarid, with a minimum age of 55.8 Ma. All node age priors were set as log normal distributions, with an offset of the minimum age of the node and given a monophyletic constraint. For the crown Acanthuridae age of 48.5 Ma, we set the mean and standard deviation of the distribution to be 2.07 and 1, respectively. For dating the split of acanthurids from luvarids, we set the mean and standard deviation as 2.26 and 1, respectively. For the overall age of acanthurids plus their eupercarian outgroups we set a minimum age at 100.1 Ma, an average of the estimated ages from recent phylogenomic higher-level studies (Betancur-R et al., 2017; Hughes et al., 2018; Near & Thacker, 2024, Santaquiteria et al., 2026) with a mean and standard deviation of 1.55 and 1, respectively.

*2D lateral photographs*

We collected 230 2D lateral specimen photographs for geometric morphometrics from online museum collections, primary literature, commercial aquarium sites, and our photography (Appendix B). We sampled only adult individuals to limit allometric effects on shape changes and ensured a standard photograph position. These images span 80 different species across the 6 extant genera within Acanthuridae, with an average number of photos per species being 2.85. Once landmarked, we found that within species variation (0.4293649) was much lower than our between species variation (1.473623) for our overall morphometric dataset.

*SurgeonShape app and metric calculations*

SurgeonShape (Figure S1) imports the landmarks and semilandmarks for all fish images, enables batch processing and custom user fin ray rotations, computes a set of biomechanical traits for each fish, and provides an output trait matrix for the user. SurgeonShape establishes a horizontal axis for each specimen, rotates each fish to the horizontal axis, rotates the jaws closed, rotates collapsed dorsal and anal fins to a minimum expanded position, rotates pectoral fins to 45 degrees to the horizontal and pelvic spines to -45 degrees, and provides user tools to correct artifacts in dorsoventral caudal fin position, caudal fin spread, and pectoral fin spread. However, rotation in the app cannot mitigate heavy distortion, such as twisted or bent fins. This app can compute all fin areas, fin aspect ratios (ARs), elongation ratio, pectoral fin position, and scalpel spine position. SurgeonShape is a Mac app that is freely available for download (https://github.com/mwestneat/SurgeonShape) with the present data set preloaded for exploration of soft tissue standardization tools. Extended details of the app and its functions are available in the readme on Github.

Pectoral and caudal fin ARs were calculated for each specimen in SurgeonShape using the fin landmarks and the generalized equation:

$$AR = \frac{b^2}{A}$$

where b is the maximum span of the respective fin (lines $b_{pec}$ and $b_{caudal}$ in Figure 1) and A is the surface area of the respective fin (curves 5-8 for the pectoral fin and curves and 9-11 for the caudal fin in Figure 1) . A straight line between the proximal ends of curves 9 and 11 was used to “close” the area of the caudal fin. Elongation ratio was also calculated by taking the total length of the fish and dividing it by body depth, computed as the vertical distance between the highest point on the body at the center of the dorsal fin and the lowest body point at the origin of the pelvic fin. Species-average fin ARs and elongation ratios were then calculated.

*Landmark subsets*

From our overall landmark dataset, we produced four subsets: body shape (15 fixed landmarks and 60 sliding semilandmarks along four curves), head shape (11 fixed landmarks and 38 sliding semilandmarks along two curves), pectoral fin shape (four fixed landmarks and 28 sliding semilandmarks along four curves), and caudal fin shape (six fixed landmarks and 61 sliding semilandmarks along three curves). We elected to use a large number of semilandmarks to aid in the sliding process downstream as increased numbers of semilandmarks help retain the true biological shape of the structure during data analyses. Many of these semilandmarks were removed before final analyses, so that these "helper points" would not prevent us from achieving appropriate statistical power or drown out shape variation from fixed landmarks (Zelditch et al., 2012). For the head shape landmark subset (pink line, Figure 1), every fourth sliding semilandmark was kept for curve 1, and every other sliding semilandmark was kept for curve 2, resulting in a total of 10 sliding semilandmarks across two curves. The same semilandmarks were removed for curves 1 and 2 for the body shape landmark subset (purple line, Figure 1), in addition to removing every other semilandmark for curves 3 and 4, resulting in a total of 20 sliding semilandmarks across four curves. For the caudal fin subset (green line, Figure 1), every other helper point was removed across all curves (9-11), resulting in a total of 29 sliding semilandmarks across three curves. We did not subsample the pectoral fin curve (yellow line, Figure 1). Statistics were only ran on these subsets. No statistics related to associations with variables of interest were done on our full shape data, as the number of variables (146, stemming from the x and y coordinates of our 73 landmarks after subsampling) exceeded our number of individuals (80 species shapes).

**Supplementary Results**

*Acanthuridae phylogenetic relationships*

Among the outgroups, the general percomorph topology is similar to other recent large-scale trees (Betancur-R et al., 2017). We find strong support for a sister-group relationship between the Chaetodontidae and Pomacanthidae, which form the sister group to the acanthuriform clade of *Luvarus*, *Zanclus*, plus Acanthuridae. The root node of a monophyletic Acanthuridae was resolved with high support, estimated to be about 54.6 Ma in our phylogenetic chronogram (Figure 2). Our phylogenetic tree resolves the genus *Naso* as a strongly supported monophyletic group, originating 19.5 Ma, forming the sister group to the rest of the Acanthuridae. No single *Naso* species was resolved as sister to the rest of the genus. Rather, three *Naso* clades were identified. The first group, *N. minor* and sisters *N. caeruleocauda* and *N. thynnoides*, is strongly supported and is first to split from the rest of *Naso* (15.5 Ma). A second group of *Naso* species, including two sister groups 1) *N. unicornis* + *N. tergus* and *N. lituratus* + *N. elegans* as well as 2) *N. hexacanthus* + *N. caesius* and *N. vlamingii* + *N. brevirostris*, were strongly supported as the overall sister group to the clade containing *N. maculatus*, *N. annulatus*, *N. lopezi*, *N. brachycentron*, *N. mcdadei* + *N. reticulatus*, *N. fageni*, and *N. tonganus* + *N. tuberosus*. Almost all nodes are heavily supported in this clade except for *N. fageni*'s sister relationship to *N. tonganus* + *N. tuberosus* (posterior probability: 0.83) and this overall clade's sister relationship to *N. mcdadei* + *N. reticulatus* (0.76). Slightly younger than *Naso*, the genus *Prionurus* (16 Ma) was also resolved as a monophyletic group with strong support and the second group to branch from the backbone topology of the Acanthuridae. All nodes within this clade are heavily supported, with *Pr. microlepidotus* recovered as the sister to the rest of the genus, *Pr. biafraensis* and *Pr. laticlavius* being sister to the clade containing *Pr. maculatus* and the sister pair *Pr. chrysurus* + *Pr. scalprum*.

At about 43 Ma, the rest of the acanthurids are split into two clades with strong support, one containing the monotypic genus *Paracanthurus* and genus *Zebrasoma*, and the other containing all *Acanthurus* and *Ctenochaetus* species . All nodes are heavily supported in the first group, with *Pa. hepatus* splitting from *Zebrasoma* at about 29 Ma. Within the genus, *Z. desjardinii* and *Z. velifer* form a clade that is sister to the rest of the group. A sister relationship is also recovered between *Z. flavescens* and *Z. scopas*. The second group containing the genera *Acanthurus* and *Ctenochaetus* is the largest but has the most weakly supported internal nodes compared to other clades within the phylogeny. Additionally, both *Acanthurus* and *Ctenochaetus* are not monophyletic genera. At about 28 Ma, two clades form, one containing 14 species of *Acanthurus*, and the other containing the rest of the *Acanthurus* species, with *Ctenochaetus* being nested within. In the clade containing just *Acanthurus* species, two sister groups split at 27 Ma, although this split is not well supported (0.7). The first clade contains *A. lineatus* + *A. sohal* and its weakly supported sister clade (0.77) comprising of *A. nigrofuscus, A. leucopareius*, and *A. nigroris* + *A. nigros*, and the other contains *A. polyzona* + *A. triostegus*, weakly supported as sister (0.5) to the clades of *A. coeruleus* + *A. guttatus*, and *A. achilles, A. leucosternon*, *A. japonicus* + *A. nigricans*.

At nearly 27 Ma, *A. thompsoni* splits from the second group containing the rest of *Acanthurus* and *Ctenochaetus*, although this split is not well supported (0.7). Two groups then split at 25.3 Ma, one containing *A. pyroferus* + *A. tristis* as a sister pair to the clade comprising of all *Ctenochaetus* species and *A. nubilus*, and the other containing the remaining 19 *Acanthurus* species. *C. striatus* and *C. hawaiiensis* + *C. tominiensis* form a paraphyletic group to the rest of the *Ctenochaetus* species and their sister taxa *A. nubilus* (weakly supported: 0.51). Within this clade, the only weakly supported node is the sister relationship between *C. flavicauda* and *C. truncatus* (0.64). The 19 remaining *Acanthurus* species in the second group from two clades at nearly 22 Ma. All relationships in the first are well supported, with *A. tennentii* and *A. olivaceus* + *A. reversus* being sister to *A. monroviae*, *A. chirurgus*, and *A. bahianus* + *A. tractus*. The second group is composed of *A. nigricauda*, *A. auranticavus* + *A. leucocheilus* (all nodes well supported) sister to *A. bariene* + *A. dussumieri*, *A. albimento*, *A. grammoptilus*, *A. gahhm* + *A. xanthopterus*, *A. mata*, and *A. blochii* + *A. maculiceps.* Both splits of *A. albimento* from *A. grammoptilus*, *A. gahhm* + *A. xanthopterus*, *A. mata*, and *A. blochii* + *A. maculiceps* as well as *A. gahhm* + *A. xanthopterus* from *A. mata* and *A. blochii* + *A. maculiceps* are not well supported (0.57 and 0.56 respectively). Additionally, the split of *A. grammoptilus* from *A. gahhm* + *A. xanthopterus*, *A. mata*, and *A. blochii* + *A. maculiceps* is more supported than the previously mentioned splits, but still not strongly supported (0.66). Here is where our nearly identical maximum likelihood tree (Figure S2) and the time-calibrated tree (Figure 2) differ in topology. In our maximum likelihood tree, *A. albimento* is not sister to the group of *A. grammoptilus*, *A. gahhm* + *A. xanthopterus*, *A. mata*, and *A. blochii* + *A. maculiceps*, but is instead sister to *A. bariene* + *A. dussumieri*. These taxa, *A. albimento*, *A. bariene* + *A. dussumieri,* form the sister group to *A. grammoptilus*, *A. gahhm* + *A. xanthopterus*, *A. mata*, and *A. blochii* + *A. maculiceps*.

### Supplementary Discussion

*Revised phylogenetic relationships of the Acanthuridae*

Our new surgeonfish phylogeny recovered similar relationships among the major subclades of surgeonfishes as in previously constructed trees (Clements et al., 2003; Guiasu &

Winterbottom, 1993; Holcroft & Wiley, 2008; Klanten et al., 2004; Sorenson et al., 2013; Tang et al., 1999; Tyler et al., 1989; Winterbottom, 1993). *Naso* is monophyletic and sister group to the rest of the surgeonfishes, followed by genus-level clades along the backbone of the tree formed by *Prionurus, Paracanthurus, Zebrasoma*, and then the combined *Acanthurus* + *Ctenochaetus* crown clade. Most surgeonfish genera are monophyletic, apart from *Acanthurus* and *Ctenochaetus*, and we reaffirm the conclusion originally reached by Randall (1955) and supported by Sorenson et al. (2013) that the Acanthuridae can be divided into two subfamilies, the Nasinae (consisting of the genus *Naso*), and the Acanthurinae (represented by the rest of the family). Within the subfamily Acanthurinae, *Prionurus* is sister to the rest of the subfamily. *Paracanthurus* is sister to *Zebrasoma*, and that clade is sister to the crown clade of *Acanthurus* and *Ctenochaetus*. Within this crown clade, we support the *Ctenochaetus* and *Acanthurus* paraphylies as shown in many other studies (Clements et al., 2003; Klanten et al., 2004; Sorenson et al., 2013).

While these relationships remain the same, we see that node ages of these clades differ. While present hypothesis indicates a nearly identical crown age of all Acanthuridae (54.58 Ma) to the 54 Ma of the Sorenson et al. (2013) topology, we estimate other major crown ages to be older. The crown ages of Nasinae (19.45 Ma), Acanthurinae (47.14 Ma), and *Prionurus* (15.99 Ma) were all older in our analysis than the Sorenson et al. (2013) topology (17 Ma, 42 Ma, and 13 Ma, respectively) but did fall within the previous authors' 95% highest posterior density (HPD) range. This is not the case for the crown ages of the clade containing *Paracanthurus* and *Zebrasoma* and the clade containing *Acanthurus* and *Ctenochaetus*. We estimate these crown ages to be 28.9 Ma and 27.66 Ma respectively, which fall outside the 95% HPD interval of the Sorenson et al. (2013) topology and are quite older than the previous authors' reported crown ages of 17 Ma and 21 Ma respectively.

Looking within each major subclade, we have mostly improved, strong nodal support, with the clade containing *Acanthurus* and *Ctenochaetus* having the highest concentration of weakly supported nodes. Relationships within the clades *Prionurus* and *Paracanthurus* + *Zebrasoma* are congruent with Sorenson et al. (2013), although we do have additional taxa that deepen our understanding of the relationships withing those subclades. We also differ from the most recent topology of the surgeonfishes in terms of the relationships within other subclades. For example, with strong support, we do not recover *N. lopezi* as the sister to all other *Naso* species. In fact, we find 1) *N. lopezi* nested within the genus, serving as the sister to a smaller clade containing *N. tonganus* + *N. tuberosus*, *N. fageni, N. reticulatus* + *N. mcdadei,* and *N. brachycentron* and 2) no single *Naso* species being sister to the rest of the genus. Instead, we find the clade of *N. minor* and *N. caeruleocauda* + *N. thynnoides*, a clade Sorenson et al. (2013) found nested further within *Naso*, to be the sister clade to the rest of the genus. More species-level differences exist in the *Naso* genus between this tree and earlier ones. For example, we resolve *N. reticulatus* and *N. mcdadei* as sisters, as opposed to *N. reticulatus* and *N. maculatus* (Sorenson et al., 2013). Additionally, *N. brevirostris* and *N. vlamingii* form a group sister to *N. caesius* + *N. hexacanthus*, as opposed to a paraphyly (Sorenson et al., 2013).

Further differences between our topology and more recent topologies are seen in the larger *Acanthurus* + *Ctenochaetus* clade. Strikingly, *A. thompsoni* and *A. triostegus* were not found to be outgroups to *Ctenochaetus* and the rest of the *Acanthurus* species. Instead, they are found nested within separate clades, although 1) *A. thompsoni* is the outgroup to a smaller subclade within the larger *Acanthurus* + *Ctenochaetus* clade and 2) nodes that determine these relationships are weakly supported. We don't support any single taxa serving as the outgroup to this larger *Acanthurus* + *Ctenochaetus* clade. Additionally, while we recover a similar tree topology constructed by Sorenson et al. (2013) in which *A. pyroferus* and *A. nubilus* both appear as members of a larger clade containing all *Ctenochaetus* species, there are differences as well. In our hypothesis, the clade of *Ctenochaetus* species that *A. nubilus* is the outgroup to does not contain *C. tominiensis*; instead, *C. tominiensis* (and its sister *C. hawaiiensis*) are strongly supported as the outgroup to a clade of *Ctenochaetus* species and *A. nubilus*. Most other differences in tree topology between our hypotheses and previous work were minor or are the result of the addition of more species to the phylogeny.

### Supplementary Figures

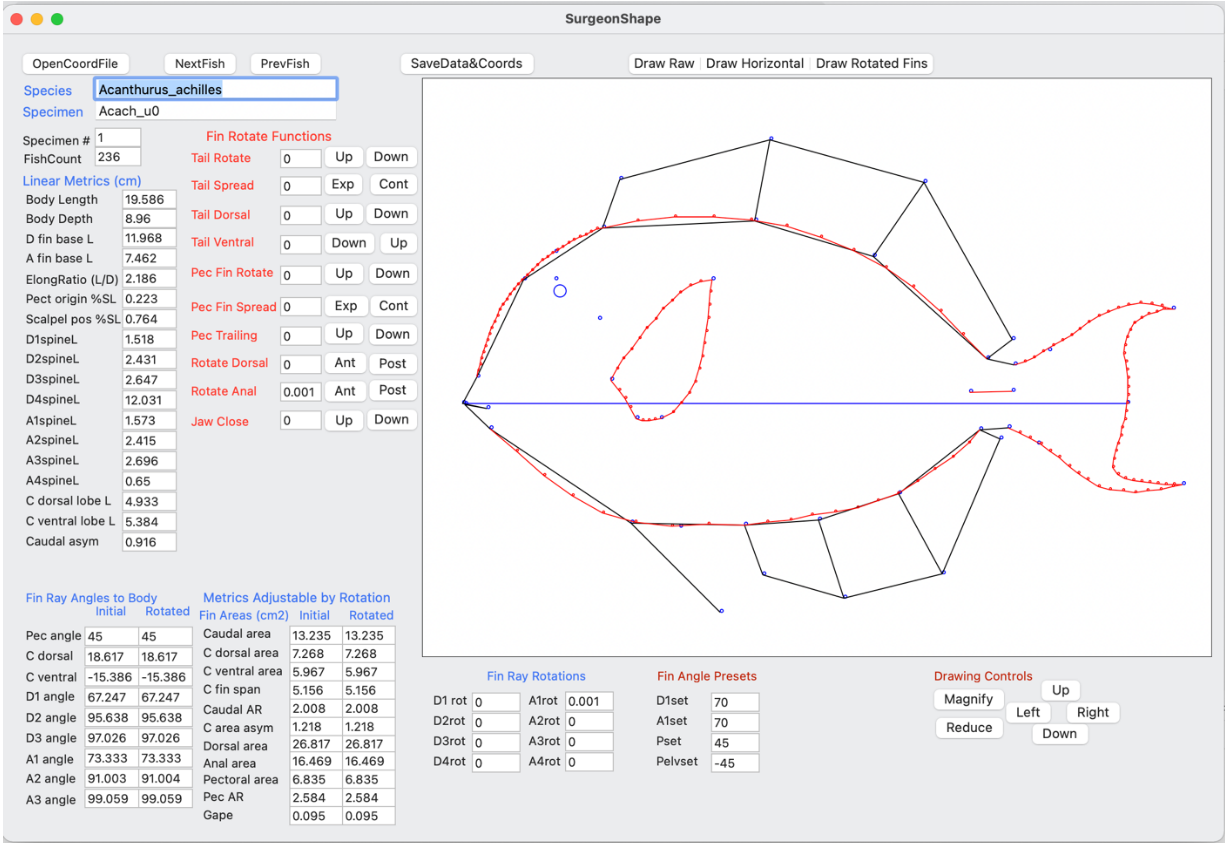


Figure S1. **SurgeonShape App.** User interface of the SurgeonShape app for the rotation of fin rays, standardization of fin angles and image artifacts, and computation of fin areas and biomechanical metrics of surgeonfish shapes. Landmarks and the horizontal fish axis are in blue, semilandmarks exported as equidistant coordinates from geomorph curves are in red, and outline and fin ray line segments drawn by the app are in black.

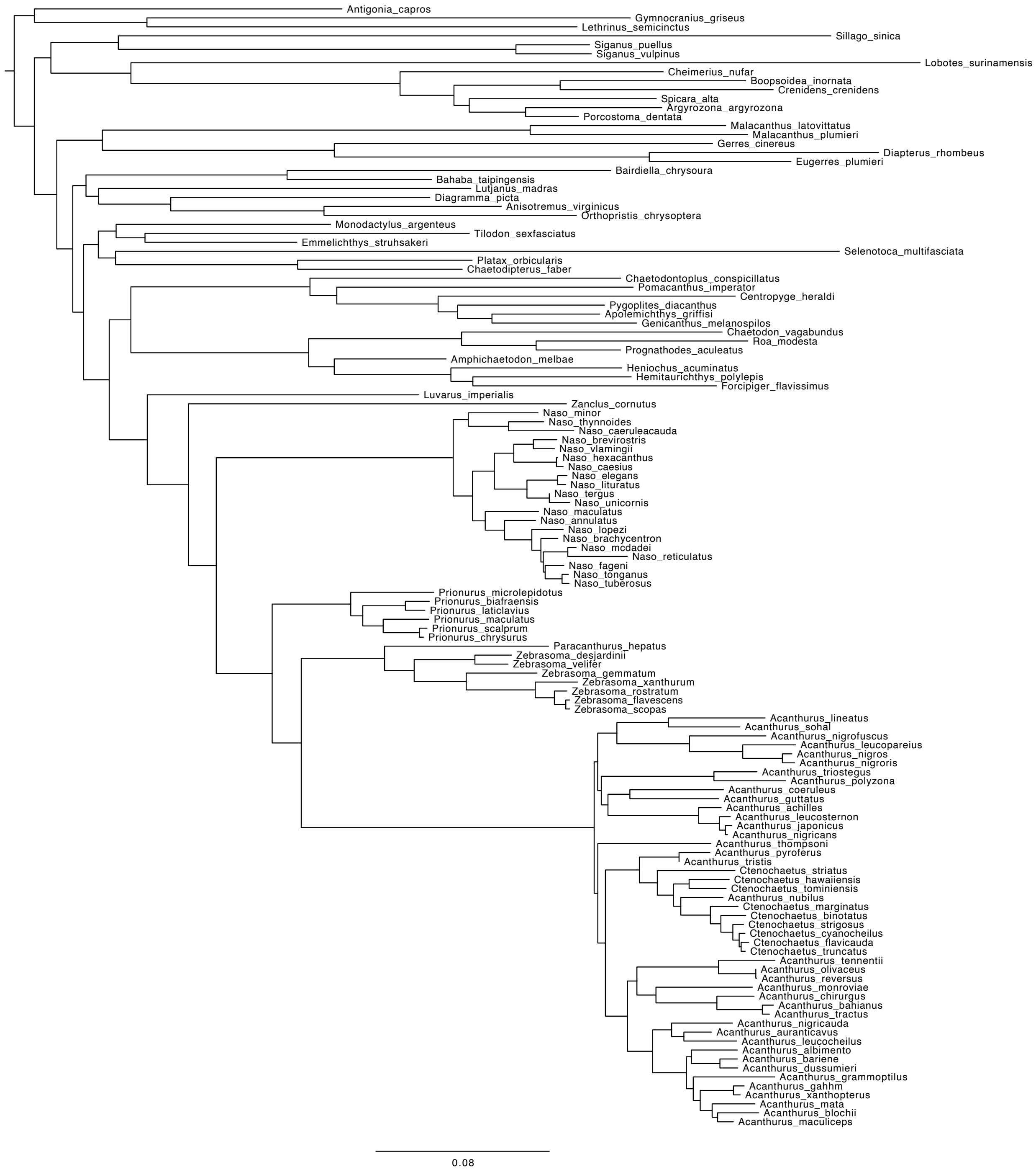


Figure S2. **Maximum-likelihood Tree for Acanthuridae.** Phylogenetic hypothesis for the Acanthuridae and outgroups based on maximum likelihood via IQ-TREE.

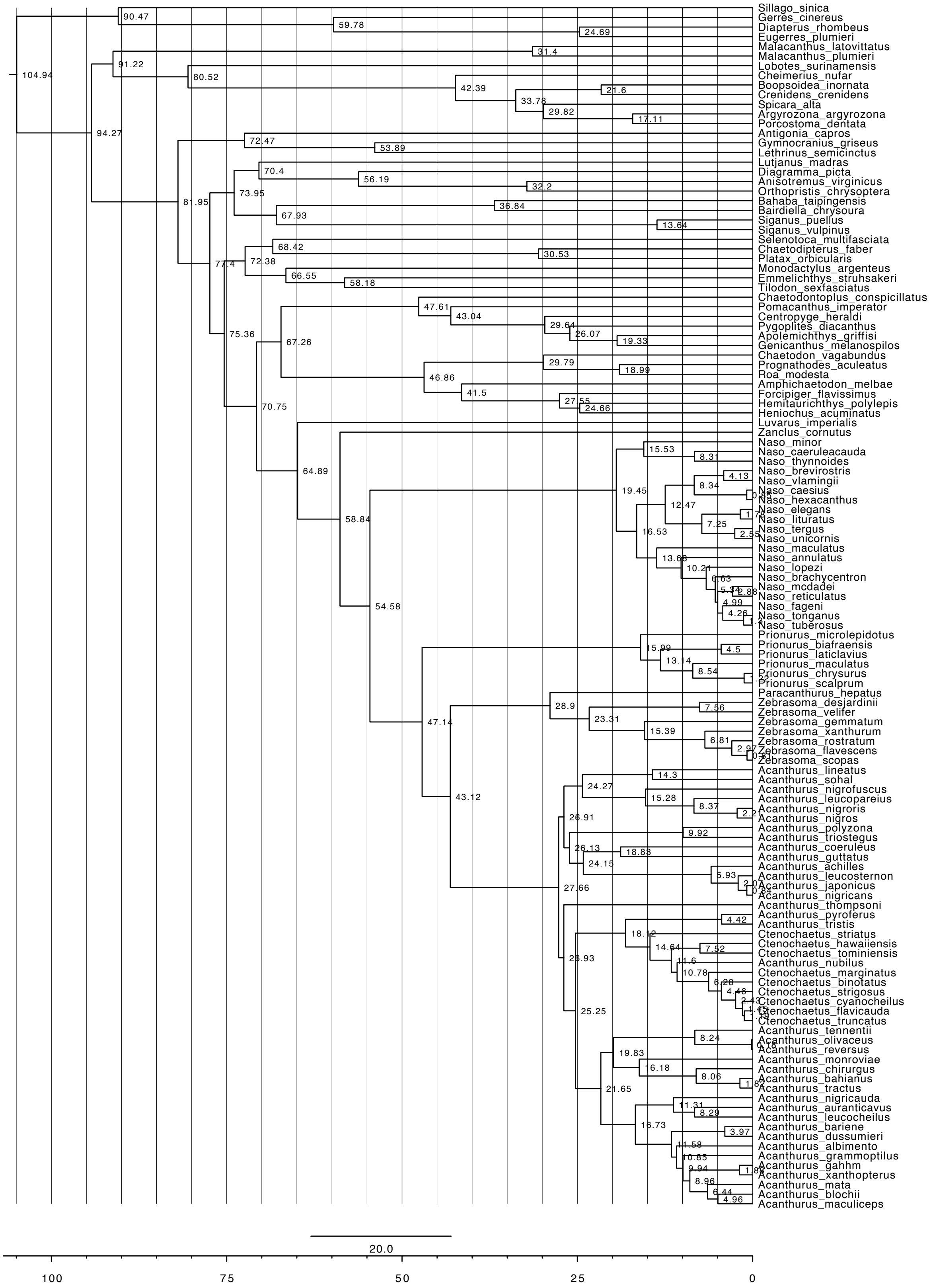


Figure S3. **Estimated Ages for All Nodes in New Phylogeny.** Node ages listed on the time-calibrated phylogenetic hypothesis of the Acanthuridae with the inclusion of all outgroups.

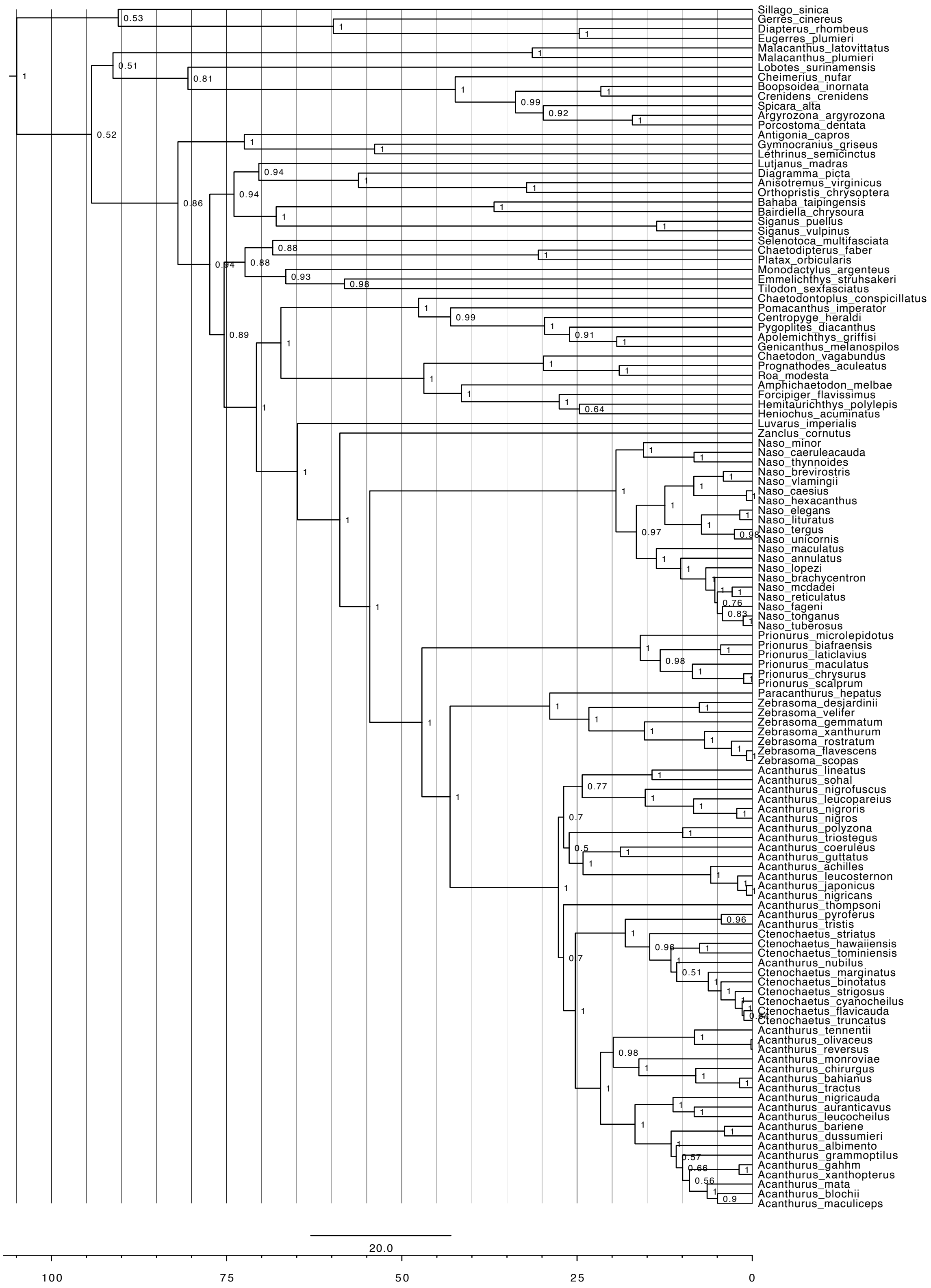


Figure S4. **Posterior Probabilities for All Nodes in New Phylogeny.** Phylogenetic hypothesis from Bayesian analysis, depicted as a majority rule consensus tree of the 95% credible set of trees with support values at nodes.

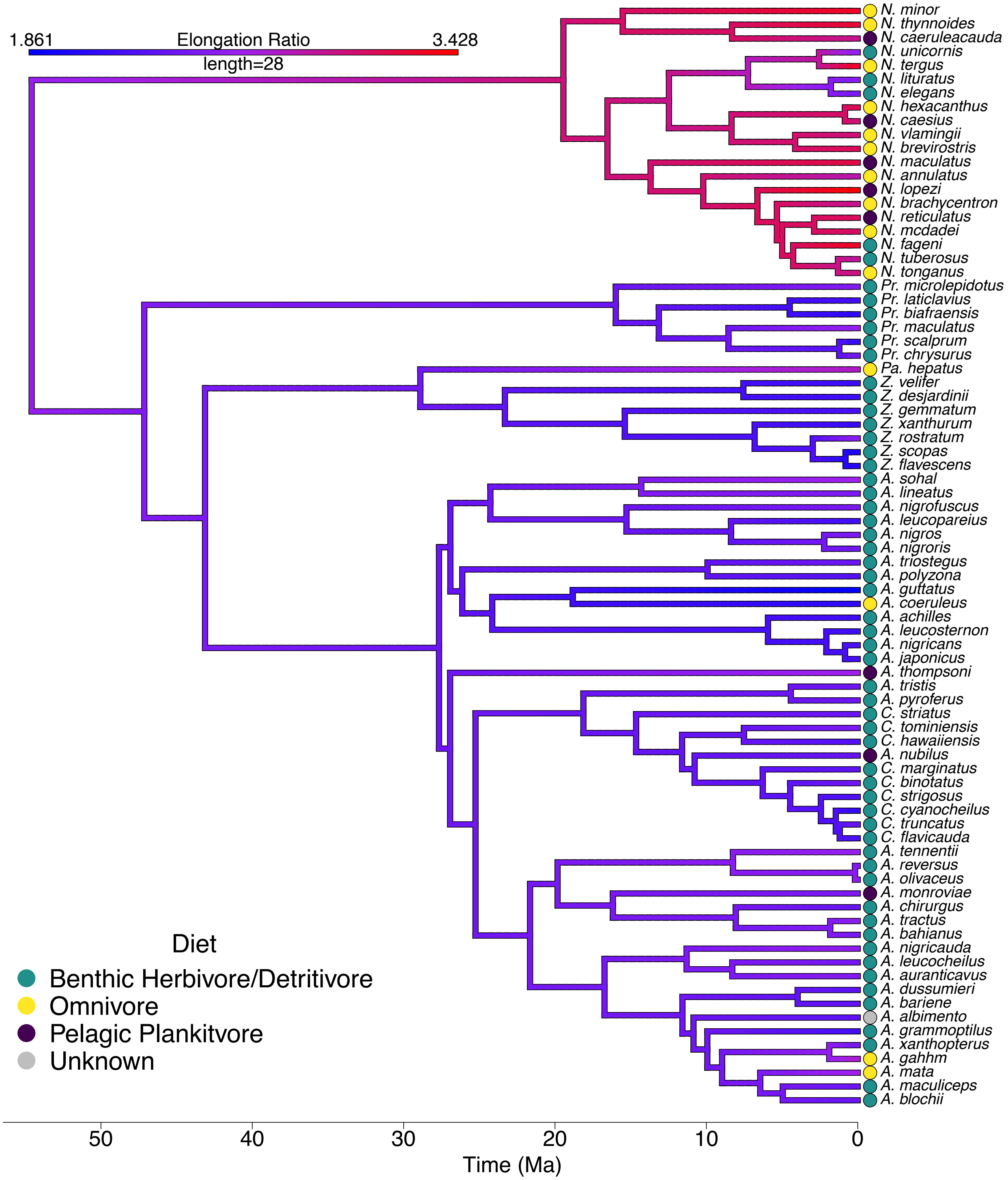


Figure S5. **Elongation Ratio along Acanthuridae Phylogeny.** Elongation ratio mapped along the Acanthuridae phylogeny by estimating states at internal nodes using maximum likelihood, with ratio values ranging from 1.861 to 3.428. Length is measured in Ma, and time is also denoted at the bottom scale. Dietary ecotype for each species is represented by colored dots at the tips of trees. Most instances of pelagic planktivory and omnivory as well as high elongation ratios are observed in the *Naso* genus.

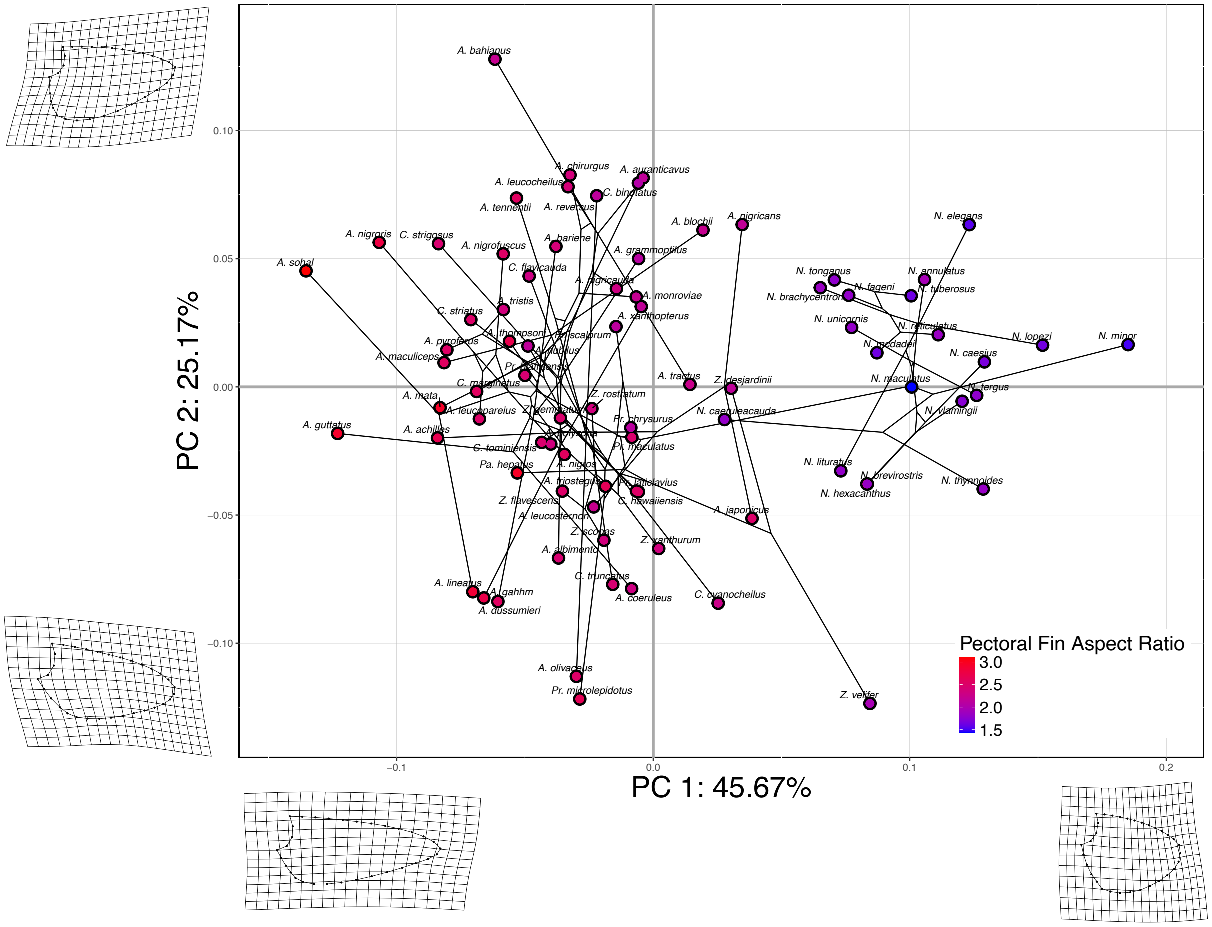


Figure S6. **Pectoral Fin Phylomorphospace.** PCA plot showing PC1 (45.67%) vs PC2 (25.17%) of the 80 shapes representing the pectoral fin shape of each species in the dataset. Points are color-coded on a gradient of pectoral fin AR. Cooler colors represent lower values of pectoral fin AR whereas warmer colors represent higher values. Thin plate splines of the minimum and maximum values of each PC are shown at the corresponding ends of each axis as a deformation of the mean shape of all 80 specimens.

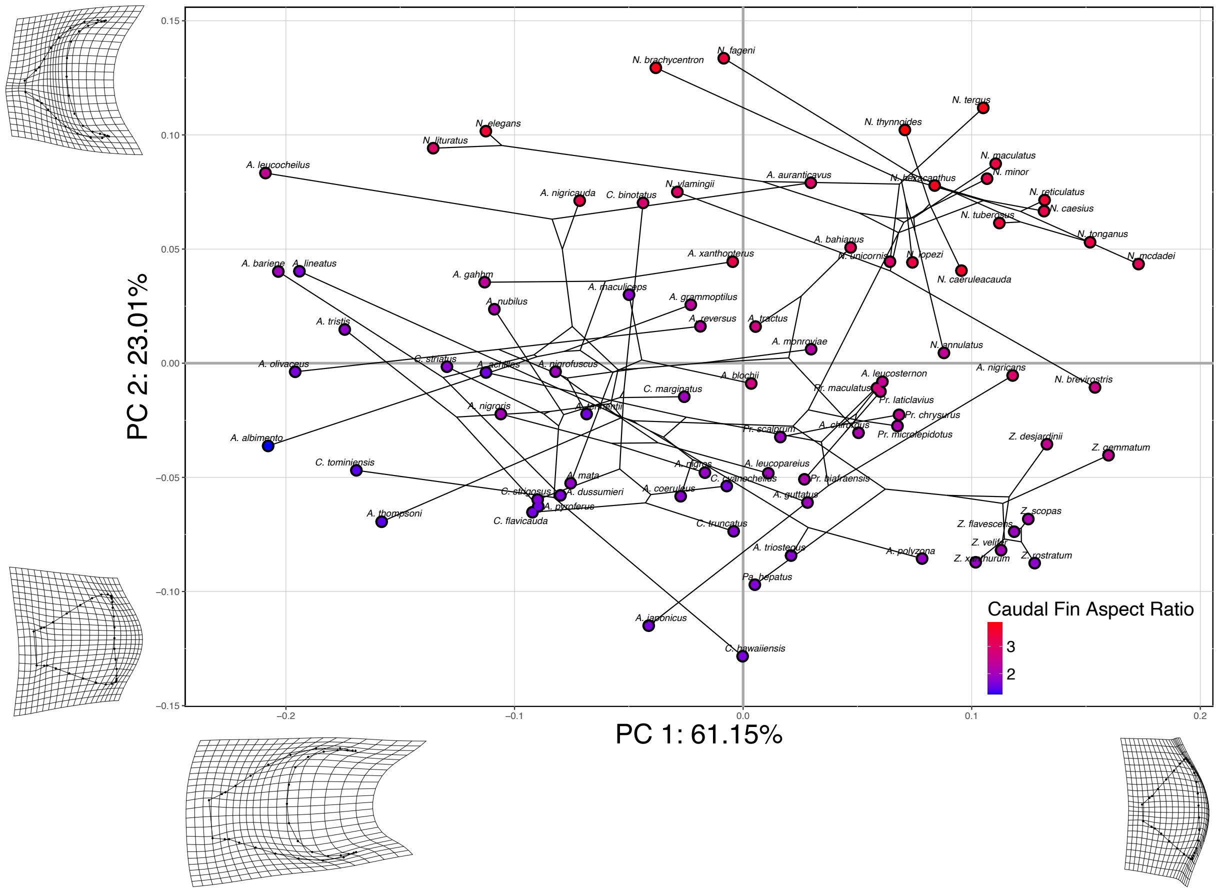


Figure S7. **Caudal Fin Phylomorphospace.** PCA plot showing PC1 (61.15%) vs PC2 (23.01%) of 79 shapes representing the caudal fin shape of each species in the dataset, with *A. sohal* removed as it is an outlier. Points are color-coded on a gradient of caudal fin AR. Cooler colors represent lower values of caudal fin AR whereas warmer colors represent higher values. Thin plate splines of the minimum and maximum values of each PC are shown at the corresponding ends of each axis as a deformation of the mean shape of all 79 specimens.

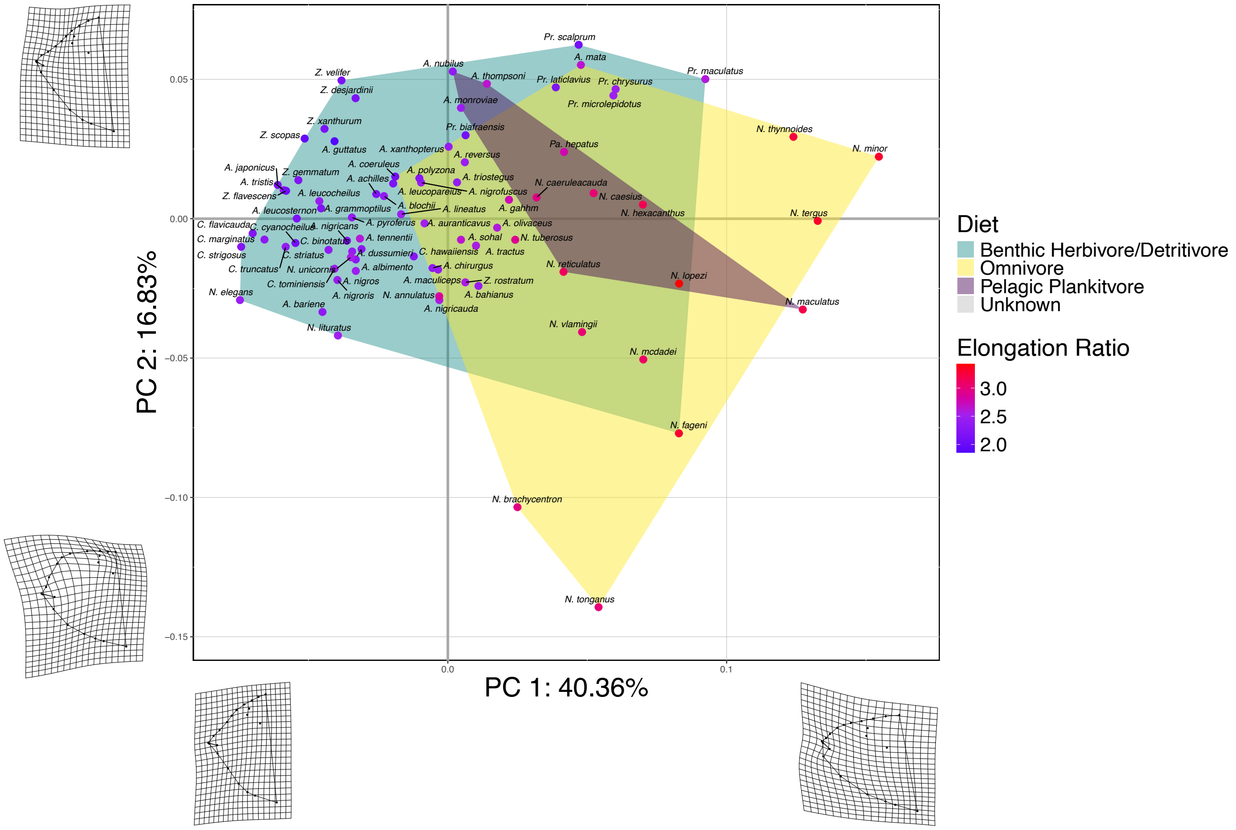


Figure S8. **Head Shape Morphospace.** PCA plot showing PC1 (40.36%) vs PC2 (16.83%) of 79 shapes representing the head shape of each species in the dataset, with *N. brevirostris* removed as it is an outlier. Dietary ecotypes are indicated by convex hulls and points are color-coded on a gradient to indicate the species mean of elongation ratio. Cooler colors represent lower values of elongation ratio whereas warmer colors represent higher values of elongation ratio. Thin plate splines of the minimum and maximum values of each PC are shown at the corresponding ends of each axis as a deformation of the mean shape of all 79 specimens.

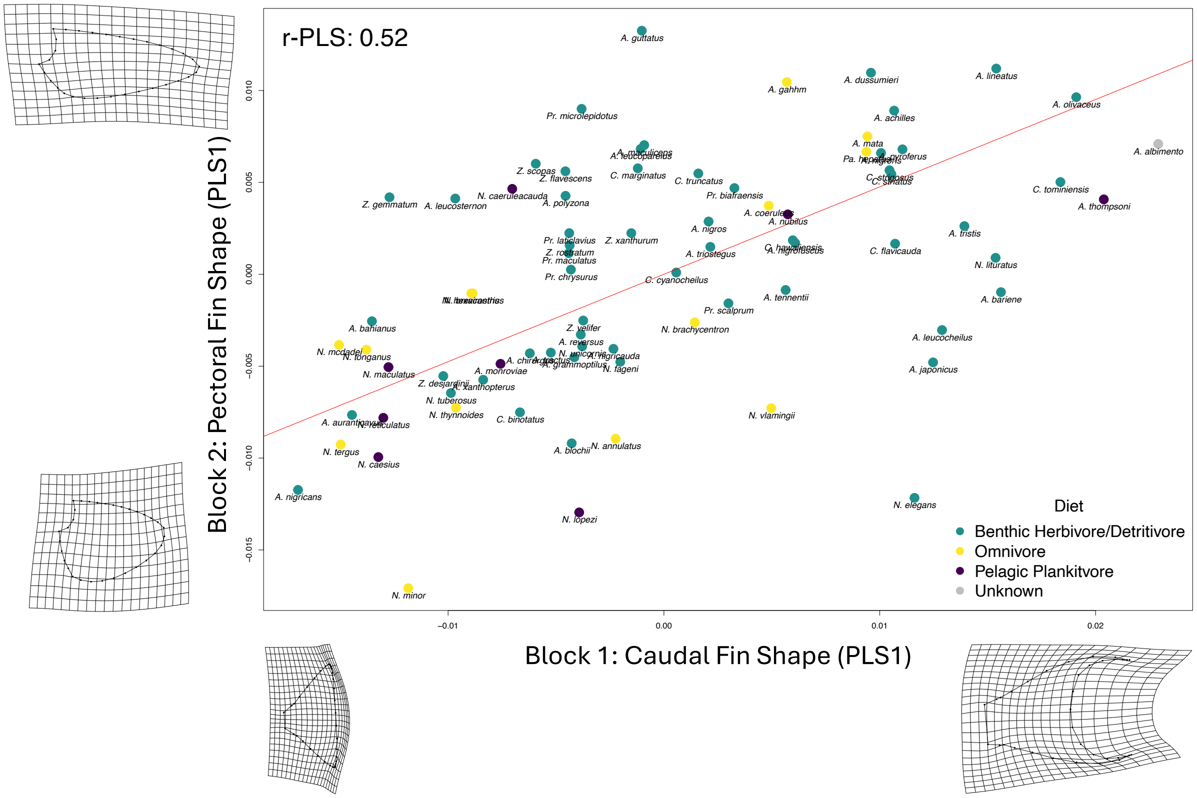


Figure S9. **Caudal Fin Shape vs Pectoral Fin Shape 2B-PLS.** Plot showing the first PLS scores for caudal fin shape (block 1) and pectoral fin shape (block 2). Caudal fin shape and pectoral fin shape are significantly paired (p-value < 0.01), with a PLS correlation coefficient of 0.52. The red line indicates the correlation between the two PLS axes yielded by a phylogenetically informed PLS test. Thin plate splines at axes' extremes visualize shape change across their respective axes. Points are color-coded based on dietary ecotype.

**Table S1. Substitution Models.** The best fit partitioning scheme determined by IQ-TREE 2. Unique: number of unique site patterns. Infor: number of parsimony-informative sites. Invar: number of invariant sites. Const: number of constant sites (can be subset of invariant sites). Site Sub Model: site substitution model. LogL; log likelihood. AIC, w-AIC: Akaike Information Criterion scores and weights. AICc, w-AICc: corrected AIC scores and weights. BIC, w-BIC: Bayesian information criterion scores and weights.

| **Gene** | **Seq** | **Site** | **Unique** | **Infor** | **Invar** | **Const** | **Site Sub Model** | **LogL** | **AIC + w-AIC** | **AICc + w-AICc** | **BIC + w-BIC** |
|---|---|---|---|---|---|---|---|---|---|---|---|
| 12S + 16S | 115 | 1501 | 876 | 616 | 727 | 727 | GTR + F + R5 | -24454.846 | 48943.692 + 0 | 48944.105 + 5.09e-322 | 49034.028 + 2.35e-314 |
| 4c4 + rag2 + rag1 | 112 | 2762 | 1428 | 918 | 1463 | 1463 | TIM2e + I + R3 | -24745.632 | 49509.264 + 0 | 49509.330 + 5.09e-322 | 49562.578 + 2.35e-314 |
| ND3 + ND5 + CytB | 101 | 3306 | 1926 | 1706 | 1409 | 1409 | GTR + F + I + R6 | -86505.118 | 173050.236 + 0 | 173050.492 + 5.09e-322 | 173172.306 + 2.35e-314 |
| COI | 122 | 652 | 322 | 269 | 367 | 367 | GTR + F + I + G4 | -18881.81 | 37785.621 + 0 | 37786.033 + 5.09e-322 | 37834.901 + 2.35e-314 |
| Otx + Zic1 | 113 | 1494 | 299 | 131 | 1255 | 1255 | HKY + F + R3 | -4797.706 | 9613.411 + 0 | 9613.532 + 5.09e-322 | 9661.194 + 2.35e-314 |
| BMP4 + DLX + MYH6 + PLAGL2 + Rhod | 98 | 3331 | 1197 | 683 | 2347 | 2347 | TNe + R3 | -17960.541 | 35935.083 + 0 | 35935.117 + 5.09e-322 | 35977.860 + 2.35e-314 |
| ETS2 | 49 | 479 | 310 | 198 | 203 | 203 | GTR + F + I + G4 | -4549.952 | 9121.904 + 0 | 9122.469 + 5.09e-322 | 9167.793 + 2.35e-314 |
| ND4 + ND6 | 86 | 2144 | 1346 | 1211 | 755 | 755 | TIM2 + F + I + R5 | -55387.833 | 110807.666 + 0 | 110807.922 + 5.09e-322 | 110898.393 + 2.35e-314 |

**Table S2. Landmarks.** List of landmark descriptions used for geometric morphometrics.

| **Landmarks** |
|---|
| Center of Lower Lip |
| Center of Upper Lip |
| Dorsal Fin Anterior Base |
| Tip of First Dorsal Ray |
| Dorsal Fin Posterior Base |
| Tip of Last Dorsal Ray |
| End of Dorsal Caudal Peduncle |
| Start of Dorsal Caudal Fin Membrane |
| Dorsal Tip of Caudal Fin |
| Ventral Tip of Caudal Fin |
| Start of Ventral Caudal Fin Membrane |
| End of Ventral Caudal Peduncle |
| Anal Fin Posterior Base |
| Tip of Last Anal Ray |
| Anal Fin Anterior Base |
| Tip of First Anal Ray |
| Pelvic Fin Anterior Base |
| Pelvic Fin Anterior Tip |
| Posterior Lip Joint |
| Center of Eye |
| Pectoral Fin Dorsal Base |
| Pectoral Fin Dorsal Tip |
| Pectoral Fin Ventral Tip |
| Pectoral Fin Ventral Base |
| Start of Caudal Spine |
| End of Caudal Spine |
| Last Dorsal Fin Spiny Ray |
| Last Anal Fin Spiny Ray |
| Dorsal Fin Midpoint |
| Anal Fin Midpoint |
| Start of Forehead |
| Dorsal Eye |
| Dorsal Gill Plate |
| Ventral Gill Plate |

**Table S3. Curves.** List of curves used for geometric morphometrics including the starting and ending fixed landmark of each curve.

| Curve | Starting Landmark | Ending Landmark |
|---|---|---|
| Dorsal Pectoral Fin | Pectoral Fin Dorsal Base | Pectoral Fin Dorsal Tip |
| Distal Pectoral Fin | Pectoral Fin Dorsal Tip | Pectoral Fin Ventral Tip |
| Ventral Pectoral Fin | Pectoral Fin Ventral Tip | Pectoral Fin Ventral Base |
| Proximal Pectoral Fin | Pectoral Fin Ventral Base | Pectoral Fin Dorsal Base |
| Dorsal Caudal Fin | End of Dorsal Caudal Peduncle | Dorsal Tip of Caudal Fin |
| Center Caudal Fin | Dorsal Tip of Caudal Fin | Ventral Tip of Caudal Fin |
| Ventral Caudal Fin | Ventral Tip of Caudal Fin | End of Ventral Caudal Peduncle |
| Forehead | Start of Forehead | Dorsal Fin Anterior Base |
| Dorsal Body | Dorsal Fin Anterior Base | Dorsal Fin Posterior Base |
| Dorsal Ventral Body | Anal Fin Anterior Base | Anal Fin Posterior Base |
| Anterior Ventral Body | Ventral Gill Plate | Anal Fin Anterior Base |

**Table S4**. **2B-PLS Results across Methodologies**. Summary of the results from 2B-PLS tests done on each morphological subset pairing (except head shape vs body shape) for tests operating under the current "all or nothing assumptions" ($\lambda$ =1 and $\lambda$ =0, respectively) as well as our Pagel's λ adjustment. Raw p-values and Holm-Bonferroni-corrected p-values are stated, along with the PLS correlation coefficients (r-PLS) and effect sizes (Z) for each comparison. (**p < 0.01, *p < 0.05)

| **Block 1 Shape** | **Block 2 Shape** | **p-value (raw)** | **p-value (corrected)** | **r-PLS** | **Effect Size (Z)** |
|---|---|---|---|---|---|
| *Pagel's λ adjustment* | | | | | |
| Caudal | Pectoral | 0.001** | 0.005** | 0.522 | 3.4304 |
| Caudal | Head | 0.027* | 0.081 | 0.431 | 1.9684 |
| Pectoral | Head | 0.051 | 0.102 | 0.446 | 1.6347 |
| Caudal | Body | 0.001** | 0.005** | 0.588 | 3.6737 |
| Pectoral | Body | 0.194 | 0.194 | 0.335 | 0.903 |
| *$\lambda = 0$* | | | | | |
| Caudal | Pectoral | 0.001** | 0.005** | 0.707 | 5.714 |
| Caudal | Head | 0.001** | 0.005** | 0.621 | 4.5946 |
| Pectoral | Head | 0.001** | 0.005** | 0.639 | 4.1666 |
| Caudal | Body | 0.001** | 0.005** | 0.725 | 4.1474 |
| Pectoral | Body | 0.001** | 0.005** | 0.762 | 4.9226 |
| *$\lambda = 1$* | | | | | |
| Caudal | Pectoral | 0.001** | 0.005** | 0.751 | 3.0293 |
| Caudal | Head | 0.029* | 0.046* | 0.471 | 1.8861 |
| Pectoral | Head | 0.023* | 0.046* | 0.506 | 1.9982 |
| Caudal | Body | 0.001** | 0.005** | 0.596 | 3.2153 |
| Pectoral | Body | 0.001** | 0.005** | 0.603 | 2.9053 |
